# Tissue-resident skeletal muscle macrophages promote recovery from viral pneumonia-induced sarcopenia in normal aging

**DOI:** 10.1101/2025.01.09.631996

**Authors:** Constance E. Runyan, Lucy Luo, Lynn C. Welch, Ziyan Lu, Fei Chen, Maxwell J. Schleck, Radmila A. Nafikova, Rogan A. Grant, Raul Piseaux Aillon, Karolina J. Senkow, Elsie G. Bunyan, William T. Plodzeen, Hiam Abdala-Valencia, Craig Weiss, Laura A. Dada, Edward B. Thorp, Jacob I. Sznajder, Navdeep S. Chandel, Alexander V. Misharin, G.R. Scott Budinger

**Affiliations:** Division of Pulmonary and Critical Care Medicine, Department of Medicine, Simpson Querrey Lung Institute for Translational Sciences. Northwestern University. Chicago, IL, USA; Department of Neuroscience, Northwestern University Feinberg School of Medicine. Chicago, IL, USA; Department of Pathology, Northwestern University Feinberg School of Medicine, Chicago, IL, USA

**Author notes:** These authors contributed equally (Alexander V. Misharin, G.R. Scott Budinger).

## Abstract

Sarcopenia, which diminishes lifespan and healthspan in the elderly, is commonly exacerbated by viral pneumonia, including influenza and COVID-19. In a study of influenza A pneumonia in mice, young mice fully recovered from sarcopenia, while older mice did not. We identified a population of tissue-resident skeletal muscle macrophages that form a spatial niche with satellite cells and myofibers in young mice but are lost with age. Mice with a gain-of-function mutation in the Mertk receptor maintained this macrophage-myofiber interaction during aging and fully recovered from influenza-induced sarcopenia. In contrast, deletion of Mertk in macrophages or loss of Cx3cr1 disrupted this niche, preventing muscle regeneration. Heterochronic parabiosis did not restore the niche in old mice. These findings suggest that age-related loss of Mertk in muscle tissue-resident macrophages disrupts the cellular signaling necessary for muscle regeneration after viral pneumonia, offering a potential target to mitigate sarcopenia in aging.

## Introduction

Sarcopenia is clinically recognized as a loss of skeletal muscle mass and function and is frequently observed in the elderly as part of normal aging^1^. Sarcopenia can also be caused or exacerbated by disease, disuse, and malnutrition^2^. Even before the COVID-19 pandemic, worsened sarcopenia was recognized as a common sequela of pneumonia, which is the most common cause of death from an infectious disease worldwide, and which disproportionately affects the elderly^3,4^. Since 2019, nearly half a billion people have been infected with severe acute respiratory syndrome coronavirus-2 (SARS-CoV-2) and more than 6 million have died from SARS-CoV-2 pneumonia over the course of the Coronavirus Disease-2019 (COVID-19) pandemic^5^. Some of the billions who survived their infection have developed dysfunction across organ systems that persists after the viral infection has cleared. These various symptoms define the post-acute sequelae of COVID-19 (PASC) syndrome and include worsening sarcopenia^6,7^.

Elegant experiments in murine models have investigated the mechanisms of impaired skeletal muscle recovery in old mice after acute damage to the muscle by toxins or thermal injury^8^. In these models, muscle regeneration is accomplished by the proliferation of satellite cells, a partially-committed progenitor cell population that forms a discontinuous layer of cells distributed along the length of the muscle fiber between the myofiber membrane and the basal lamina^9^. In young adult mice satellite cells proliferate after direct skeletal muscle injury and fuse with myofibers to regenerate muscle, but this process fails in old mice^10–12^. This failure results from cell-autonomous dysfunction in the satellite cells^13–16^ combined with a loss of signals to muscle satellite cells from the niche, including Notch ligands^10,11^, FGFs^17,18^, TGF-β, Wnt ligands^19,20^, activators of p8 MAPK pathways^21–24^, and inhibitors of JAK/STAT signaling^25,26^, among others^27,28^. These signals have been suggested to originate from soluble factors in the circulation^29,30^, monocyte-derived macrophages^31,32^, regulatory T cells^33,34^ and fibroadipogenic progenitors (FAPs), a specialized fibroblast population in the muscle^35,36^.

Direct injury to the skeletal muscle results in massive necrosis of myofibers accompanied by infiltration of large numbers of inflammatory cells and soluble factors from the circulation. Infiltrating monocyte-derived macrophages have been reported to be important for skeletal muscle repair after direct injury, although the precise mechanisms are not known^31,37–41^. Direct injury differs from the skeletal muscle pathology observed in critically ill patients where infiltration by inflammatory cells is uncommon^42^. Instead, biopsies from these patients show a loss of myofiber cross-sectional area accompanied by biochemical evidence of enhanced protein degradation by the muscle-specific E3 ubiquitin ligases FBXO32 (also known as ATROGIN-1) and TRIM63 (also known as MURF1)^42^. We showed that many of these features are reproduced in mice following pulmonary infection with a murine-adapted influenza A virus^43^. Using this model, we reported that IAV infection caused sarcopenia in mice^43^. Young adult mice completely resolved their IAV-induced sarcopenia within 2 weeks after infection, but old mice never fully recovered, instead exhibiting a step decline in muscle function that persisted until their death^44^.

MERTK is a tyrosine kinase expressed on the surface of macrophages and other cells^45^. In macrophages, MERTK interacts with phosphatidylserine residues on adjacent cells and plays a critical role in phagocytosis. We reported young adult mice globally-deficient in *Mertk* failed to recover from IAV-induced sarcopenia, phenocopying findings in old mice^44^. Like many genetic interventions in young mice that mimic age-related phenotypes, these findings show the global loss of *Mertk* is sufficient for the failure to resolve IAV-induced sarcopenia but provide no information as to whether *Mertk* is necessary to resolve IAV-induced sarcopenia during normal aging. Here, we report that a loss of MERTK in tissue-resident skeletal muscle macrophages with advancing age is necessary for some features of age-related frailty in normal mice and for the failure of old mice to recover from influenza A-induced sarcopenia.

## Results

### Expression of cleavage-resistant MERTK restores skeletal muscle recovery after viral pneumonia in old mice

We previously reported that the expression of *Mertk* is reduced in flow-sorted skeletal muscle macrophages from old (18-26 months of age) compared to young adult (4-6 months of age) mice. We queried data from a recently published human skeletal muscle single cell RNA-sequencing atlas for the expression of *Mertk* in skeletal muscle macrophages as a function of age^46^. We found the levels of *Mertk* in skeletal muscle macrophages were reduced in old compared with younger individuals (**Supplemental Figure S1A**). To determine whether the loss of *Mertk* was necessary for the failure to recover from viral pneumonia in old animals, we aged a cohort of mice expressing cleavage-resistant MERTK (*Mertk*^Δ*483–488*^ hereafter referred to as *Mertk(CR*)) to 18–26 months of age. *Mertk(CR)* mice carry a mutation of *Mertk* in the proteolytic cleavage site that is targeted by extracellular protease ADAM17 to inactivate it^47^. Thus, this mutation results in a gain of MERTK function. As controls, we aged colonies of wild-type mice and mice deficient in *Mertk* (*Mertk*^-/-^) and compared them with young adult wild-type mice (4–6 months of age) (**Figure 1A and Supplemental Figure S1B**).

**Figure 1.**
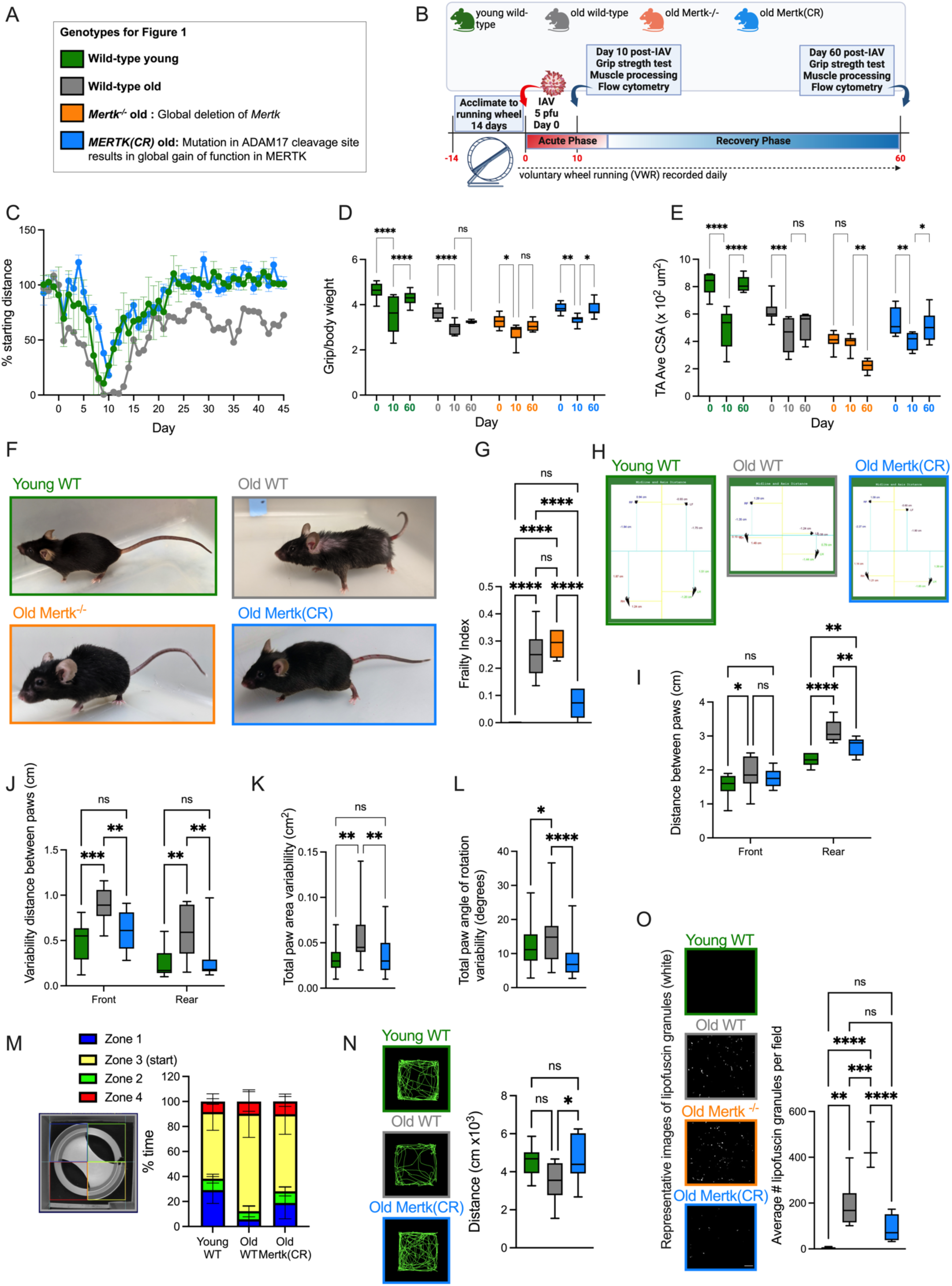
*Mertk* is necessary for the failed recovery from influenza A induced sarcopenia in old mice. (A) Murine strains and ages used in Figure 1. (B) Schematic of the experimental design. See also Figure S1C, D. (C) Daily voluntary wheel running distance before and after IAV infection (n=9-12 mice per group). See also Figure S1E and S1L. (D) Box plots of forelimb grip strength/body weight after 0 (naïve), 10- or 60-days influenza A infection in young adult wild-type, old wild-type, old *Mertk^-/-^*, and old *Mertk(CR)* mice (n = 9–12 mice per group). Two-way ANOVA was used to determine statistical significance. See also Figure S1M. (E) Box plots of average transverse cross-sectional area of tibialis anterior muscle (TA) after 0 (naïve), 10- or 60-days after influenza A infection in young adult wild-type, old wild-type, old *Mertk^-/-^*, and old *Mertk(CR)* mice (n=5-12 mice per group). Two-way ANOVA was used to determine statistical significance. See also Figure S1N, O. (F) Representative photographs of 4-month-old wild type (WT) mouse (green outline), 22-month-old wild-type (WT) (grey outline), 21-month-old *Mertk^-/-^* mouse (orange outline), and 22-month-old *Mertk(CR)* mouse (blue outline). See also video S1-3. (G) Box plots of blinded measures of frailty index in young adult wild-type, old wild-type, old *Mertk^-/-^*, and old *Mertk(CR)* mice. (n=10 mice/group). Ordinary one-way ANOVA used to determine statistical significance. (H) Representative images of front and rear paw placement and angle relative to midline for young adult wild-type, old wild-type, and old *Mertk(CR)* mice. (I) Box plots of average paw-to-paw distance (stance) in cm for front and rear paws for young adult wild-type, old wild-type, and old *Mertk(CR)* mice. (n=10-12 mice per group). Two-way ANOVA was used to determine statistical significance. (J) Box plots show average step-to-step stride width variability in front and rear limbs for young adult wild-type, old wild-type, and old *Mertk(CR)* mice (n=10-12 mice per group). Two-way ANOVA was used to determine statistical significance. (K) Box plots show the variability in maximal area of each paw placed on treadmill per step for young adult wild-type, old wild-type, and old *Mertk(CR)* mice (n=10-12 mice per group). Ordinary one-way ANOVA used to determine statistical significance. (L) Box plots show variability in angle of paw placement relative to midline, averaged for all paws per mouse for young adult wild-type, old wild-type, and old *Mertk(CR)* mice (n=10-12 mice per group). Ordinary one-way ANOVA used to determine statistical significance. (M) Image demonstrates zones of an elevated zero maze indicating exposed and covered zones and starting placement. Bar graph represents average percent of time each mouse spent in each zone over a period of 600 seconds for young adult wild-type, old wild-type, and old *Mertk(CR)* mice (n=10-12 mice per group) (N) Representative diagrams of movement tracked for 600 seconds in open field analysis for young adult wild-type, old wild-type, and old *Mertk(CR)* mice. Box plots represent average total distance traveled (n=10-12 mice per group). Ordinary one-way ANOVA used to determine statistical significance. (O) Representative images of auto-fluorescent lipofuscin granules in unstained tibialis anterior muscle cross sections (scale bar = 30mm), and box plots representing average number of lipofuscin granules per field for young adult wild-type, old wild-type, old *Mertk^-/-^*, and old *Mertk(CR)* mice (n=5-12 mice per group). Ordinary one-way ANOVA used to determine statistical significance.

To measure muscle function in mice, we offered them access to monitored running wheels, which mice use voluntarily, running several miles per day^48^. We infected running wheel-adapted young adult and old mice with IAV (A/WSN/33 [H1N1]) to induce viral pneumonia and assessed them during recovery (**Figure 1B**). Because the LD_50_ for IAV is about 4-fold lower in old compared with young adult mice^49^, we administered IAV doses titrated to induce similar mortality and weight loss in young adult and old mice (**Supplemental Figure S1C,D**). Young adult wild-type mice and old *Mertk(CR)* mice recovered pre-infection running wheel distance after IAV infection, while old wild-type mice had persistent reductions 60 days after infection (**Figure 1C and Supplemental Figure S1E**). We previously reported that young adult mice lacking the muscle-specific E3 ligase *Fbxo32* did not develop sarcopenia after influenza A infection^43^. The induction of FBXO32 was similar in young and old wild-type mice and in *Mertk(CR)* and *Mertk*^-/-^ mice (**Supplemental Figure S1F,G**), confirming that IAV-induced sarcopenia was similar across ages and genotypes. Similarly, measures of forelimb grip strength and muscle fiber diameter fell in all mice after viral infection, recovered completely in young adult wild-type mice and old *Mertk(CR)* mice, but did not recover in old wild-type or old *Mertk*^-/-^ mice (**Figure 1E**,**Figure 1F and Supplemental Figure S1H-K**).

Even before infection with influenza, old *Mertk(CR)* mice were visibly different from old wild-type mice, appearing more like young adult than old wild-type mice (**Figure 1F and Movie S1-3**). Accordingly, we measured frailty using a 31-parameter clinical frailty index^50^. Frailty scores of old *Mertk(CR)* mice were lower than old wild-type mice and similar to young adult wild-type mice (**Figure 1G**). Gait analysis suggested protection from age-related widening of rear stance, and step-to-step variability in stance width and paw placement (**Figure 1H-L**). Old *Mertk(CR)* mice showed improved exploration in zero maze **(Figure 1M)** and movement in open field (**Figure 1N**) compared to old wild-type mice. Compared with old wild-type mice, histologic examination of skeletal muscle from old *Mertk(CR)* mice showed reduced accumulation of highly-autofluorescent lipofuscin, while old *Mertk*^-/-^ mice showed increased accumulation of lipofuscin (**Figure 1O**). Nevertheless, compared to young adult wild-type mice, running wheel distance, grip strength, cross sectional area of the tibialis anterior muscle and weight of the soleus and extensor digitorum longus muscles were similarly reduced in old wild-type, *Mertk^-/-^*and *Mertk(CR)* mice (**Supplemental Figure S1L-Q**). Collectively, these data suggest that maintaining MERTK activity with advancing age allows recovery from influenza A-induced sarcopenia in old animals and confers resistance to some measures of muscle fitness and movement without affecting age-related sarcopenia.

### Tissue-resident skeletal muscle macrophage Mertk and Cx3cr1 are necessary for muscle satellite cell expansion during recovery from influenza A pneumonia-induced sarcopenia

We performed a flow cytometry analysis of single-cell suspensions generated from skeletal muscle tissue during recovery from influenza A pneumonia-induced sarcopenia (experimental design and representative gating strategy **Supplemental Figure S2A-C**). Consistent with our previous report, a population of tissue-resident skeletal muscle macrophages is expanded between 4–6 days after influenza A infection in young adult mice and persists for at least 10 days after infection (**Supplemental Figure S2E**). This expansion of tissue-resident skeletal muscle macrophages is accompanied by an expansion of muscle satellite cells and FAPs in young adult mice (**Supplemental Figure S2F,G**). In old mice, we observed no expansion of tissue-resident skeletal muscle macrophages, muscle satellite cells, or FAPs during recovery from IAV pneumonia (**Supplemental Figure S2E-G**). In contrast, the expansion of tissue-resident skeletal muscle macrophages, muscle satellite cells, and FAPs after IAV infection was similar in old *Mertk(CR)* mice compared to young adult wild-type mice 10 and 60 days after viral pneumonia (**Figure 2B,C and Supplemental Figure S2H**). Consistent with our previous reports, tissue-resident skeletal muscle macrophages were MERTK+, MHCIl^lo^ and ∼30% express the fractalkine receptor CX3CR1 (**Supplemental Figure S2I**)^44^.

**Figure 2:**
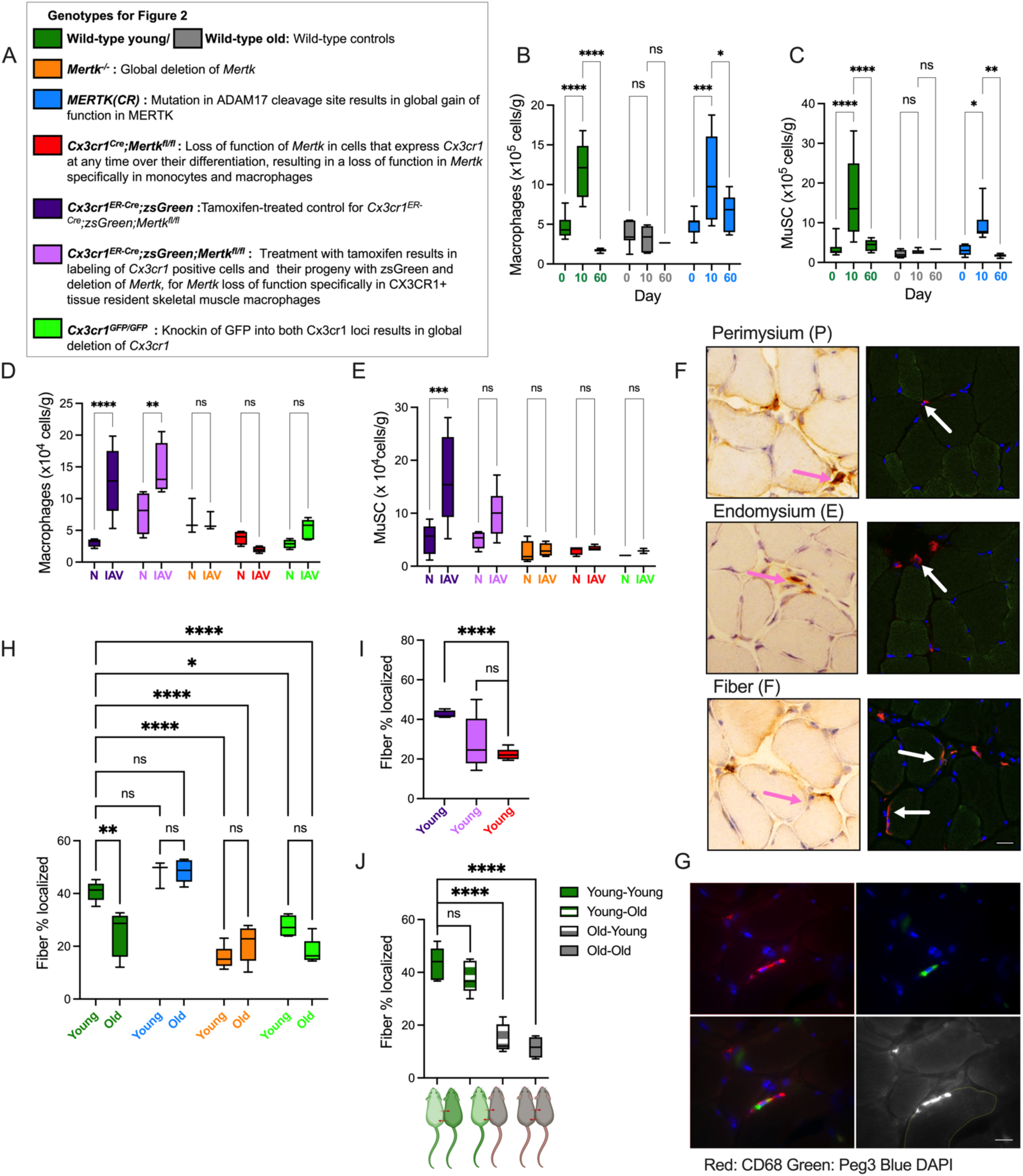
The localization and expansion of tissue resident skeletal muscle macrophages and satellite cells after influenza infection requires *Cx3cr1* and *Mertk* in young adult mice, is lost in old mice, and is maintained in old *Mertk(CR)* mice. (A) Murine genotypes used for these experiments. (B) Macrophage populations were quantified using flow cytometry from single cell suspensions of hindlimb muscles from young adult wild-type, old wild-type, and old *Mertk(CR)* mice at day 0 (naïve), 10- or 60- day post IAV. Box plots show total numbers of skeletal muscle macrophages per gram of tissue (n=6-12 mice per group). Two-way ANOVA was used to determine statistical significance. See also Figure S2A,B,E and S2H and G for FAPs. (C) Muscle satellite cell populations (MuSC) were quantified using flow cytometry from single cell suspensions of hindlimb muscles from young adult wild-type, old wild-type, and old *Mertk(CR)* mice at day 0 (naïve), 10- or 60-day post IAV. Box plots show total numbers of MuSC per gram of tissue (n=6-12 mice per group). Two-way ANOVA was used to determine statistical significance. See also Figure S2A,C,F and S2H and G for FAPs. (D) Macrophage populations were quantified using flow cytometry from single cell suspensions of hindlimb muscles from *Cx3cr1^ER-Cre/+^;zsGreen*, *Cx3cr1^ER-Cre/+^;zsGreen;Mertk^fl/fl^, Mertk^-/-^, Cx3cr1^Cre/+^;Mertk^fl/fl^* and *Cx3cr1^GFP/GFP^* mice at day 0 (naïve, N), or 10-day post IAV. Box plots show total numbers of skeletal muscle macrophages per gram of tissue (n=6-12 mice per group). Two-way ANOVA was used to determine statistical significance. See also Figure S2A, B and S2J for FAPs. (E) Muscle satellite cell populations (MuSC) were quantified using flow cytometry from single cell suspensions of hindlimb muscles from *Cx3cr1^ER-Cre/+^;zsGreen*, *Cx3cr1^ER-Cre/+^;zsGreen;Mertk^fl/fl^, Mertk^-/-^, Cx3cr1^Cre/+^;Mertk^fl/fl^* and *Cx3cr1^GFP/GFP^* mice at day 0 (naïve, N), or 10-day post IAV. Box plots show total numbers of MuSC per gram of tissue (n=6-12 mice per group). Two-way ANOVA was used to determine statistical significance. See also Figure S2A, C and S2J for FAPs. (F) Representative images of macrophage localization in either the perimysium, endomysium or abutting muscle fibers (scale bar is 20μm). See also Figure S3B-D. (G) Representative images showing fiber-associated macrophages (red) in proximity to satellite cells (green) (scale bar is 10μm). (H) Box plots display the percent of total macrophages that are localized with fibers in naïve young adult mice or old mice from wild-type, *Mertk^-/-^* and *Cx3cr1^GFP/GFP^* (n=3-5 mice per group) Two-way ANOVA was used to determine statistical significance. See also Figure S3E and F for perimysium and endomysium localized macrophages. (I) Box plots display the percentage of total skeletal muscle macrophages localized with fibers in young adult *Cx3cr1^ER-Cre;^zsGreen* and *Cx3cr1^ER-Cre;^zsGreen; Mertk^fl/fl^* mice after treatment with tamoxifen by oral gavage 10 and 9 days before harvest, and in young adult *Cx3cr1^Cre^; Mertk^fl/fl^* mice. (n=4-5 mice per group) Unpaired t test was used to determine statistical significance. See also Figure S3 G,H for perimysium and endomysium localized macrophages. (J) Young adult and old mice were joined as heterochronic parabionts for 60 days as indicated. Box plots display the percentage of total skeletal muscle macrophages localized with fibers in the individual pairs (n=3-6 mice per group).Unpaired t test was used to determine statistical significance. See also Figure S4A-C for confirmation of blood chimerism and S4E,F for perimysium and endomysium localized macrophages.

We previously reported that mice globally-deficient in *Mertk* or *Cx3cr1* failed to expand tissue-resident skeletal muscle macrophages and muscle satellite cells after viral pneumonia^44^. In that report, we used a lineage tracing system that labels circulating monocytes to show the expansion of skeletal muscle macrophages after influenza A-induced sarcopenia is not a result of recruitment of circulating monocytes and their differentiation into monocyte-derived macrophages in the skeletal muscle^44^. Because Mertk is expressed in cells other than macrophages, we generated mice lacking Mertk specifically in monocytes and macrophages (*Cx3cr1^Cre^*;*Mertk^fl/fl^*). and mice with an inducible loss of *Mertk* in *Cx3cr1*-expressing macrophages (*Cx3cr1^ER-Cre^;zsGreen;Mertk^fl/fl^*, and control *Cx3cr1^ER-Cre^;zsGreen*), which we treated with tamoxifen by oral gavage 9 and 10 days before infection with IAV and harvested them 10 days after infection with influenza A virus. We have shown using this inducible Cre recombinase system that 10 days after tamoxifen treatment >80% of circulating monocytes, which also express *Cx3cr1*, are replaced by new unlabeled monocytes from the bone marrow^51^.

We compared these strains with mice globally-deficient in Mertk (*Mertk^-/-^*) and mice globally-deficient in *Cx3cr1* (*Cx3cr1^GFP/GFP^*) in which a *GFP* knock-in prevents expression of *Cx3cr1*. Young adult mice from all of these mutant strains were infected with influenza A and muscle homogenates were analyzed using flow cytometry. The expansion of skeletal muscle macrophages, satellite cells, and FAPs was similar in wild-type young adult mice and control mice (*Cx3cr1^ER-Cre^;zsGreen*) treated with tamoxifen before influenza A infection (**Figure 2B-E and Supplemental Figure S2H,J**). Deletion of *Mertk* from all monocytes and macrophages (*Cx3cr1^Cre^*;*Mertk^fl/fl^*) prevented the proliferation of skeletal muscle macrophages, muscle satellite cells, and FAPs after influenza A infection (**Figure 2D,E and Supplemental Figure S2J**). In mice lacking *Mertk* in tissue-resident skeletal muscle macrophages expressing *Cx3cr1* (*Cx3cr1^ER-Cre^;zsGreen;Mertk^fl/fl^*treated with tamoxifen), skeletal muscle macrophages expanded after influenza A infection, but muscle satellite cells did not (**Figure 2D,E**). Skeletal muscle macrophages, satellite cells, and FAPs did not expand in mice globally lacking *Mertk* or *Cx3cr1* (**Figure 2D,E and Supplemental Figure S2J**). These results show that *Mertk* in tissue-resident skeletal muscle macrophages that express *Cx3cr1* is necessary for satellite cell proliferation after IAV pneumonia.

### MERTK preserves the localization of skeletal muscle tissue-resident macrophages with myofibers in old mice

In the developing brain, investigators discovered that microglia require *Mertk* and *Cx3cr1* to remove pieces of living neurons (synaptic pruning), a process they called “trogocytosis”^52–54^. In zebrafish, muscle satellite cell proliferation after laser-induced injury requires the removal of skeletal muscle membrane patches enriched in phosphatidylserine from myofibers by monocyte-derived macrophages^41^. We therefore wondered whether *Mertk* and *Cx3cr1* were important for the localization of skeletal muscle macrophages to the satellite cell niche and whether this niche was disrupted with advancing age. We used immunofluorescence and immunohistochemistry to estimate the steady-state localization of skeletal muscle macrophages in young adult and old wild-type mice, old *Mertk(CR)* mice, and young adult mice from the mutant strains described above.

Within tibialis anterior muscle cross sections, we observed macrophages within the perimysium and endomysium with some endomysial macrophages abutting the fibers in a wrapped configuration (**Figure 2F and Supplemental Figure S3B-D**). These fiber-associated macrophages make up ∼40% of skeletal muscle macrophages in young adult wild-type mice but are less frequently associated with skeletal muscle fibers in old mice (**Figure 2H**). The fiber-associated macrophages are localized with the muscle satellite cells at the fiber periphery (**Figure 2G**). The abundance of fiber-associated macrophages was maintained in old *Mertk(CR)* mice at a level similar to young adult wild type mice (**Figure 2H**). Fiber-associated macrophages were significantly less abundant in young adult or old *Mertk^-/-^*mice and young adult *Cx3cr1^GFP/GFP^*mice (**Figure 2H**, percentages of macrophages in the perimysium and endomysium are shown in **Supplemental Figure S3E,F**). Fiber-associated skeletal muscle macrophages were lost in young adult mice lacking *Mertk* in all monocyte and macrophage populations (*Cx3cr1^ER-Cre^;Mertk^fl/fl^*) but were maintained in young adult *Cx3cr1^ER-Cre^;zsGreen;Mertk^fl/fl^* mice 10 days after tamoxifen treatment (**Figure 2I**, percentages of macrophages in the perimysium and endomysium are shown in **Supplemental Figure S3G,H**). Heterochronic parabiosis restores defective muscle satellite cell proliferation and skeletal muscle repair after direct injury in old mice, perhaps via circulating factors like GDF11 that enhance Notch signaling^11,55^. To determine whether heterochronic parabiosis could reverse the loss of tissue-resident, fiber-associated skeletal muscle macrophages in old mice, we generated heterochronic and control parabionts and harvested them after 60 days. We analyzed blood samples using flow cytometry to confirm chimerism (**Supplemental Figure S4A-C**). We quantified the localization of skeletal muscle macrophages in tibialis anterior muscle from the exterior limbs of the parabiont pairs using immunohistochemistry. Heterochronic parabiosis did not reverse the age-related loss of muscle volume (**Supplemental Figure S4D**) nor did it restore the normal localization of skeletal muscle macrophages after injury (**Figure 2J**, percentages of macrophages in the perimysium and endomysium are shown in **Supplemental Figure S4E, F**).

### MERTK and CX3CR1 support the localization of macrophages to myoblasts where they promote their proliferation and differentiation

Our findings suggest CX3CR1 and MERTK coordinate the localization of macrophages to muscle fibers and this localization is necessary for MERTK to promote muscle satellite cell proliferation. We used an *in vitro* co-culture system to visualize the interactions of primary macrophages with muscle fibers. We cultured a C2C12 myoblast cell line either alone or with equal numbers of primary bone marrow-derived macrophages (BMDM) from wild-type or mutant mice (**Figure 3A**). C2C12 cells proliferate and fuse to form myotubes *in vitro* over the course of 7 days in the absence of macrophages (**Figure 3B)** and the expression of the only known ligand of CX3CR1, CX3CL1, is induced during myocyte fusion coincident with the increased expression of myogenin **(Supplemental Figure S5A-F**). Co-culture of BMDM from young adult or old wild-type mice or *Mertk(CR)* mice enhanced the proliferation and differentiation (measured as percent of fused nuclei) of C2C12 myoblasts when compared with no BMDM treated control cells (**Figure 3C and D**). In contrast, BMDM derived from *Mertk^-/-^* or *Cx3cr1^GFP/GFP^* mice did not enhance the proliferation or differentiation of C2C12 cells compared with no BMDM control cells (**Figure 3C, D and F**). These interactions were reciprocal; co-culture of BMDM with C2C12 cells resulted in the proliferation of BMDM from young adult and old wild-type mice and in young adult *Mertk(CR)* mice but BMDMs from mice lacking *Cx3cr1* or *Mertk* did not proliferate in response to co-culture with C2C12 cells (**Figure 3E and F**). To determine whether a direct interaction between BMDMs and C2C12 cells was necessary to promote their proliferation, we co-cultured BMDMs from young adult wild-type mice with C2C12 cells directly or separated by a semipermeable membrane in the same well (**Figure 3G**). C2C12 cells only proliferated when in direct contact with BMDM (**Figure 3H**). In contrast, BMDM still proliferated when separated from the fusing myofibers **(Figure 3I)**. These findings suggest that direct contact is required for macrophages to influence the behavior of myoblasts, but that macrophages may respond to secreted factors from the differentiating muscle cells. We previously reported that adult mice deficient in *Il6* or treated with a murine adapted antibody targeting the IL-6 receptor (tocilizumab) were protected against IAV-induced sarcopenia^43^. To model IAV induced sarcopenia *in vitro*, we allowed C2C12 myofibers to differentiate and then treated them with IL-6 or vehicle to induce fiber atrophy^43^ (**Figure 3J, K**). We then rinsed the IL-6 from the plates and allowed wild-type BMDM to interact with the fibers for 2 hours. More macrophages were associated with fibers that had been treated with IL-6 **(Figure 3L, M)**.

**Figure 3.**
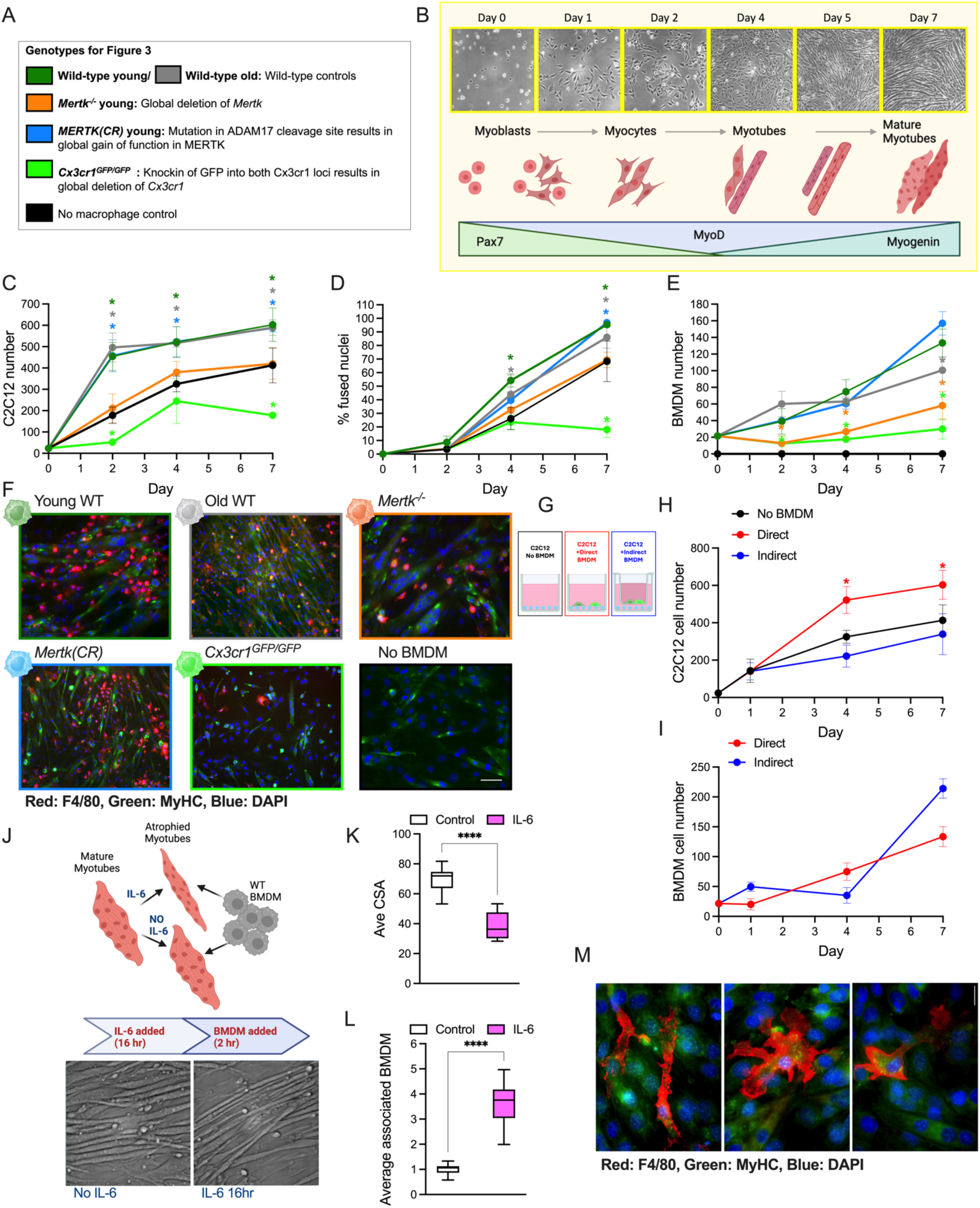
MERTK and CX3CR1 support the localization of macrophages to myoblasts where they promote their proliferation and differentiation. (A) Genotypes of mice used to generate bone marrow derived macrophages (BMDM) for this figure. (B) Top: Representative phase contrast images of C2C12 cells in 2-D culture in the 7 Days after inducing their differentiation. Bottom: Schematic demonstrating processes of myoblast proliferation and their differentiation into myocytes, marked by reduced expression of PAX7 and increased expression of MYOD followed by their fusion to form multinucleated myotubes that subsequently mature marked by increasing expression of MYOGENIN. See also Figure S5. (C) C2C12 cells were co-cultured with BMDM from young adult wild-type, old wild-type or young *Mertk(CR), Mertk^-/-^* or *Cx3cr1^GFP/GFP^* mice, or cultured without the addition of BMDMs. Cell counts derived from total nuclei counts minus the number of macrophages averaged for 3-5 fields per well of a 6-well plate, with each well representing a data point (n=3-10 wells per sample) * indicates significant difference between the genotypes compared to no macrophage control (ANOVA followed by Tukey’s comparison). (D) C2C12 cells were co-cultured with BMDM from young adult wild-type, old wild-type or young *Mertk(CR), Mertk^-/-^* or *Cx3cr1^GFP/GFP^* mice, or cultured without the addition of BMDMs. The percentage of C1C12 cells have fused nuclei (two or more nuclei per cell) is shown. 3-5 images per well of a 6-well plate were analyzed and averaged for each well (n=3-10 wells per sample) * indicates significant difference between the genotypes compared to no macrophage control (ANOVA followed by Tukey’s comparison). (E) C2C12 cells were co-cultured with BMDM from young adult wild-type, old wild-type or young *Mertk(CR), Mertk^-/-^*or *Cx3cr1^GFP/GFP^* mice. The number of BMDMs is shown. Cell counts derived from counts of F4/80+ macrophages per field averaged for 3-5 fields per well of a 6-well plate, with each well representing a data point (n=3-10 wells per sample) * indicates significant difference between the genotypes compared to BMDM from young adult wild-type mice (ANOVA followed by Tukey’s comparison). (F) Representative images of C2C12 cells cultured for 7 days with BMDMs generated from young adult wild-type, old wild-type or young *Mertk(CR), Mertk^-/-^* or *Cx3cr1^GFP/GFP^* mice, or cultured without the addition of BMDMs. Macrophages are stained with red (F4/80) and C2C12 cells with MHY3 (green) BMDMs from *Mertk^-/-^* and *Cx3cr1^GFP/GFP^* mice failed to proliferate with C2C12 cells. (G) BMDMs from young adult wild-type mice were co-cultured in direct contact with C2C12 cells or separated by a semipermeable membrane. (H) The number of C2C12 cells when cultured in direct contact with BMDMS from young adult wild type mice, separated from them by a semipermeable membrane, or cultured without BMDM. Cell counts derived from total nuclei counts minus the number of macrophages averaged for 5 fields per well of a 6-well plate, with each well representing a data point (n=3-7 wells per sample) * indicates significant difference between C2C12 cells cultured without BMDM (ANOVA followed by Tukey’s test). (I) The number of BMDMs generated from young adult mice when they were cultured in direct contact with C2C12 cells or separated from them by a semipermeable membrane. Cell counts derived from counts of F4/80+ macrophages per field averaged for 3-5 fields per well or membrane, with each well or membrane representing a data point (n=3-7 per sample). No significant differences were identified (ANOVA). (J) C2C12 cells were differentiated to myofibers and then treated with 10ng/ml recombinant murine IL-6 for 16 hours to induce atrophy followed by rinse and co-culture with media control or BMDMs from young adult wild type mice for 2 hours. Top: Schematic of experimental design. Bottom: Representative phase contrast images of C2C12 cells treated with vehicle or IL-6. (K) Measurement of myofiber average cross-sectional area 50/field x 3 fields per well of a 6well plate from C2C12 cells differentiated to myofibers followed by treatment with IL-6 for 16 hours (n=12 wells). Unpaired t test was used to determine statistical significance. (L) Average number of BMDM per field 5 fields per well. (n=9 wells). Unpaired t test was used to determine statistical significance. (M) Representative images of BMDM from wild-type young adult mice co-cultured with C2C12 cells that had been previously treated with IL-6. BMDM are stained with F4/80 (Red) and C2C12 cells are stained with MYH3 (Green). Several distinct macrophage morphologies were observed.

### Single cell RNA-sequencing suggests a signaling niche that includes skeletal muscle macrophages and satellite cells during recovery from influenza A pneumonia

We performed single cell RNA-sequencing of single cell suspensions of skeletal muscle homogenates enriched for flow-sorted macrophages, FAPs, and satellite cells from naïve or influenza day 10, young adult and old wild-type mice (**Figure 4A**). The single cell object can be explored at https://www.nupulmonary.org/. We annotated cell types using previously published single cell data sets^56,57^ (**Figure 4A, Supplemental Figure S6A**). Unsupervised clustering identified subpopulations of cells reported by other groups, all of which were represented in our data set (**Figure 4B, C, Supplemental Figure S6A**). Within our single cell RNA-sequencing data, we identified a population of skeletal muscle macrophages that express express *Mertk* and high or low levels of *Cx3cr1*(**Figure 4A,B**). These cells also express *Spp1, Trem2, Gpnmb,* and *Lgals3*, and genes with demonstrated importance in efferocytosis including Stabilin 1 (*Stab1*), T cell immunoglobulin receptor 2 (*Tim2*, *Havcr2*), *Axl*, integrin αV (*Itgav*), integrin β5 (*Itgb5*) and *Cd36.* This population has been associated with skeletal muscle dysfunction in muscle fibrosis in a murine model of muscular dystrophy^56^. Compared to other skeletal muscle macrophage populations, *Spp1*+*Cx3cr1*+ skeletal muscle macrophages expressed higher levels of genes involved in survival and migration (*Csf1r*, *Mmp14*, *Cxcr4* and *Cxcl16*) and reduced expression of inflammatory genes (*Ccl2*, *Ccl6*, *Ccl7*, *Ccl9, Tnf, Mif and IL7r*) and genes related to lipid processing (*Plin2*, *lpl*, *Fabp4* and *CD36)* (**Figure 4B**).

**Figure 4.**
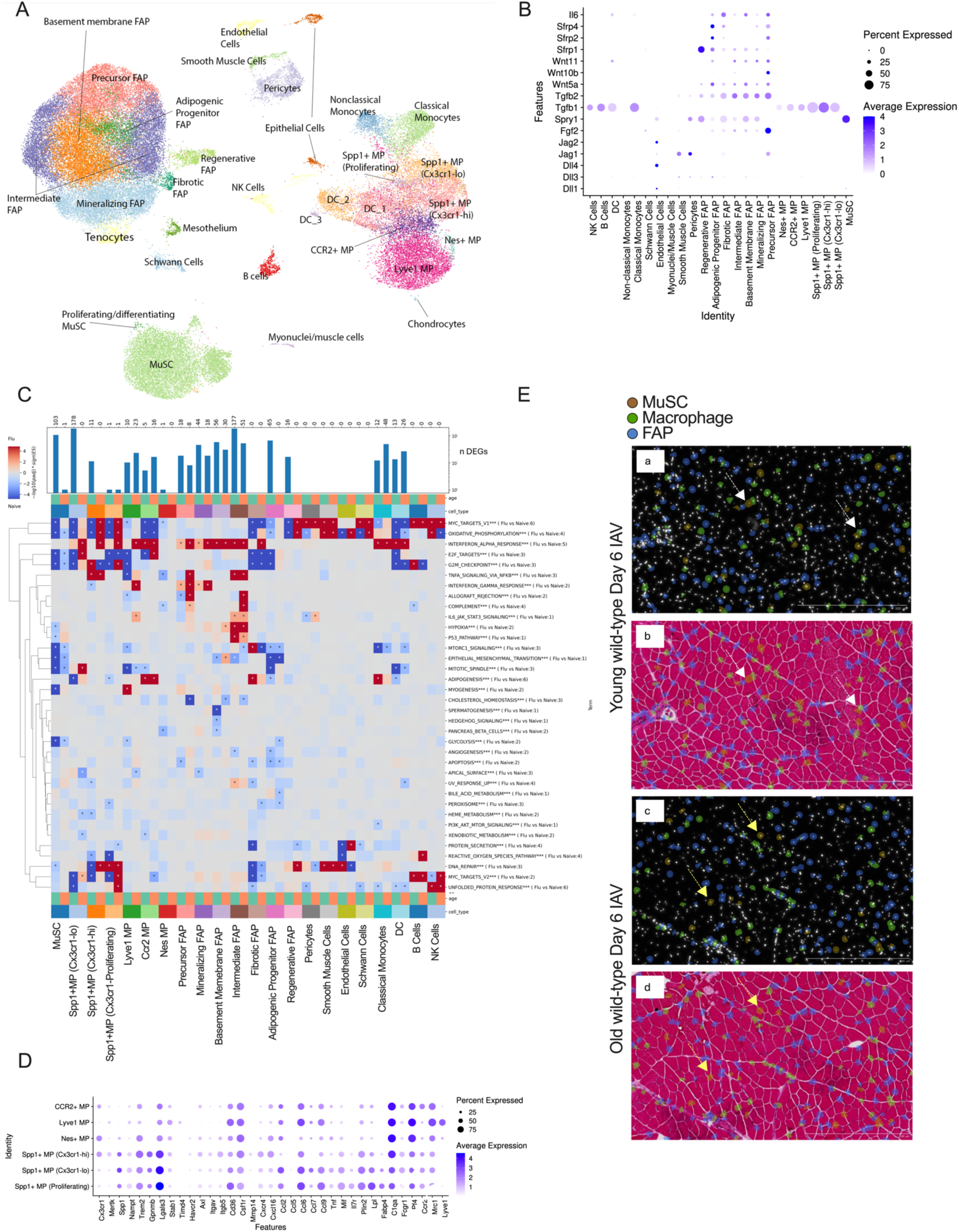
Single cell RNA-sequencing suggests a signaling niche that includes skeletal muscle macrophages and satellite cells during recovery from influenza A pneumonia. (A) Uniform manifold approximation and projection (UMAP) plot showing single-cell RNA-seq analysis of single cell suspensions of hindlimb muscles from young or old mice before or 10 days after influenza A infection after sorting to enrich for myeloid, MuSC and FAP populations. See also Figure S6A. (B) Dot plots with marker genes defining macrophage clusters. See also Figure S6A. (C) Pseudobulk differential gene expression in each cell population comparing naïve with influenza A infected mice in young adult and old mice. The number of differentially expressed genes is indicated in the bar graph above the heatmap. Heatmap indicates relative expression of genes in the indicated GSEA pathway. Only GSEA pathways that were significantly differentially expressed in one or more cell populations are shown. Significant changes are indicated with a * (FDR q<0.05). See also Figure S8 A-C. (D) Dot plot indicating expression of ligands implicated in satellite cell proliferation. See also Figure S9 A-C. (E) Spatial transcriptomic images of tibialis anterior muscle from young adult (top) and old (bottom) mice 6 days after influenza A infection. Cell populations were resolved and annotated using cell-type specific markers using nuclear segmentation. a. Young adult mouse: nuclear staining highlighting nuclei from muscle satellite cells (brown), macrophages (green) and FAPs (blue). b. Same image as (a) overlaid on hematoxylin and eosin staining from the same section. White arrows denote areas of colocalization of these three cell types. c. Old mouse: nuclear staining highlighting nuclei from muscle satellite cells (brown), macrophages (green) and FAPs (blue). d. Same image as (b) overlaid on hematoxylin and eosin staining from the same section. Yellow arrows denote areas of MuSC devoid of contact with Macrophages or FAPs. Scale bars 500μm. See also Figure S10 A,B and Supplemental Table 1.

There were no significant differences in the abundance of any cell populations in the skeletal muscle as a function of age or influenza A infection (**Supplemental Figure S6B**). We examined pseudobulk differential gene expression from naïve or day 10 post influenza A infection across skeletal muscle cell populations in young adult and old mice. Fewer differentially expressed genes were observed in young adult compared to old animals after influenza infection across all cell populations (711 genes in young adult mice compared to 224 genes in old mice) (**Figure 4C and Supplemental Figure S6C**). The union of genes in young adult and old animals was small (**Supplemental Figure S7A**). We performed GSEA analysis of the differentially expressed genes across cell populations for young adult and old mice before and after influenza A infection (**Figure 4C, Supplemental Figure S7B-D**). Consistent with a response to influenza infection, genes implicated in “interferon alpha and gamma responses”, “oxidative phosphorylation” and “TNF signaling via NF-κB” were significantly changed across multiple cell populations in both young adult and old mice (**Figure 4C, Supplemental Figure S7B-D**). In young adult mice, processes involved in cell proliferation (“MYC Targets”, “E2F Targets”, “G2M Checkpoint”) were enriched in all three populations of *Spp1*+ macrophages (Cx3cr1high, Cx3cr1low, and proliferating), *Lyve1*+ (perivascular) macrophages, muscle satellite cells, and adipogenic and fibrotic FAPs (**Figure 4C**). In old mice, these processes were simultaneously upregulated only in *Lyve1*+ macrophages and proliferating *Spp1*+ macrophages (**Figure 4C**). In young and old mice, *Spp1*+ macrophages upregulated genes implicated in “unfolded protein response”, “TNF signaling via NF-κB”, “reactive oxygen species”, “heme metabolism”, “peroxisome” and “adipogenesis” (**Figure 4C**). Muscle satellite cells in young and old mice upregulated genes involved in “mitotic signal”, “MTORC1 signaling”, “epithelial mesenchymal transition”, but only satellite cells from young mice upregulated “glycolysis” and “myogenesis” (**Figure 4C**).

Our findings suggest *Mertk* is sufficient and necessary to maintain skeletal muscle macrophages near myofibers and promote the proliferation of satellite cells after influenza A pneumonia in old mice. As many of the signaling pathways causally implicated in the regeneration of muscle after direct injury involve spatially-localized intercellular interactions, we undertook a supervised analysis of our own single cell RNA-sequencing data and a published skeletal muscle single cell atlas of murine aging to ask which cells in the skeletal muscle express ligands and receptors for these pathways^10–12,17–24,58^. These include Notch ligands^10,11^, FGFs^17,18^, TGF-β and Wnt signals^19,20^ among others^21–28^. The expression of canonical Notch ligands was limited to FAPs, which expressed *Jag1*, and endothelial cells, smooth muscle cells and pericytes, which express multiple canonical Notch ligands. (**Figure 4D and Supplemental Figure S8A**). In contrast, all of the cell populations in the dataset expressed one or more Notch receptors (**Supplemental Figure S8B**). The expression of Wnt ligands was almost exclusively limited to FAPs, all of which expressed *Wnt5a* and *Wnt11*, as well as the Wnt inhibitory ligands *Sfrp1*, *Sfrp2* and *Sfrp4* (**Figure 4D and Supplemental Figure S8A**). Precursor FAPs also expressed *Wnt2* and *Wnt10b*. Satellite cells only expressed Wnt receptors *Fzd4*, *Ror2*, and *Ryk*, while FAPs expressed a wider range of Wnt receptors and Wnt modifiers (e.g. glypicans) (**Supplemental Figure S8B**). *Fgf2* ligand expression was also limited to FAPs, while the FGF2 antagonist *Spry1* was expressed by FAPs, satellite cells, and pericytes (**Figure 4D and Supplemental Figure S8A**). FGF receptors were expressed by most cell populations in the dataset, in particular, satellite cells expressed *Fgfr1* and *Fgfr4* (**Supplemental Figure S8B**). *Il6* expression was also limited to FAPs (**Figure 4D and Supplemental Figure S8A**). Skeletal muscle macrophages expressed *Tgfb1* and a handful of other genes involved in skeletal muscle regeneration including *CD44* and *Lrp1* as well as *Adam17*, which cleaves and inactivates MERTK, but notably did not express other ligands causally linked to satellite cell proliferation (**Figure 4D and Supplemental Figure S8A**).

We were surprised that with the exception of TGF-β, which is required for tissue macrophage maintenance^59^, skeletal muscle macrophages do not express ligands causally linked to satellite cell proliferation after injury before or after influenza A infection. Instead, expression of these ligands was largely limited to FAPs and endothelial cells (**Figure 4D and Supplemental Figure S8A-C**). This might be explained if skeletal muscle macrophages remove an inhibitory signal that precludes signaling by FAPs and endothelial cells. A prediction of this model is that skeletal muscle macrophages, muscle satellite cells, FAPs and endothelial cells should be found in close proximity to one another in young mice. To confirm the presence of this niche, we performed spatial transcriptomics of skeletal muscle in young adult and old animals 6 days after influenza infection using the Xenium single-cell spatial transcriptomic platform using the Mouse Multi-Tissue standard panel (see **Supplemental Table 1** for list of probes). The spatial transcriptomic object can be explored at https://www.nupulmonary.org/. We used nuclear segmentation to cluster cells and resolved 17 cell populations (**Supplemental Figure S10A**). These included muscle satellite cells, macrophages, three populations of FAPs, three populations of myonuclei including those from slow twitch fibers, endothelial cells, pericytes, smooth muscle cells, mast cells, and cells localized to the neurovascular bundle including epithelial cells, lymphatic endothelial cells, perineural cells, adipocytes, and Schwann cells (**Supplemental Figure S10A,B**). In both young adult mice and old mice, we observed skeletal muscle macrophages adjacent to muscle satellite cells and FAPs (**Figure 4E**). Although images were only obtained from one animal for each condition, there were fewer muscle satellite cells co-localized with macrophages in young adult mice (49/94 satellite cells macrophages (52.1%) compared with old mice 13/63 (20.6%), consistent with our systematic measurements of myofiber-associated macrophages (**Figure 2H-J**). Projecting these images onto hematoxylin and eosin-stained images showed niches comprised of muscle satellite cells, macrophages, and FAPs were localized near myofibers (**Figure 4E**).

## Discussion

Elderly survivors of pneumonia, including SARS-CoV-2 pneumonia, develop worsening sarcopenia that can persist for years, increasing the risk for the compounding morbidity that limits healthspan at the end of life^3,60,61^. In a severe model of pneumonia induced by influenza A virus infection, we found that young adult and old mice develop a similar worsening of sarcopenia. While young adult mice recover from IAV-induced sarcopenia within 2 weeks, old mice never do, instead suffering a step decline in skeletal muscle function. We report that a gain of function mutation in MERTK prevented the age-related loss of skeletal muscle macrophages adjacent to myofibers. Old mice harboring a gain of function mutation in MERTK recovered from IAV-induced sarcopenia with the expansion of skeletal muscle macrophages, FAPs and satellite cells similar to young adult mice. Deletion of *Mertk* in *Cx3cr1*-expressing tissue resident skeletal muscle macrophages caused a loss of myofiber-associated macrophages in young adult mice and prevented influenza A-induced sarcopenia in old mice. These results demonstrate that *Mertk* is both necessary and sufficient for the age-related failure to recover from IAV-induced sarcopenia and highlight MERTK as a potential target for anti-aging interventions to preserve skeletal muscle function during aging.

An important strength of our study stems from our model system. Almost all studies of skeletal muscle dysfunction involve direct injury to the muscle by micropuncture, toxins, thermal injury, or genetic damage to the myofiber induced by mutations associated with muscular dystrophy^62,63^. In all these models, severe muscle necrosis induces a localized inflammatory response characterized by the recruitment of neutrophils and monocyte-derived macrophages^62,63^. Unlike these models, influenza A pneumonia in mice reproduces phenotypes reported in patients with critical illness. These include a reduction in fiber cross-sectional area accompanied by an increase in the muscle-specific E3 ubiquitin ligase ATROGIN-1 in the absence inflammatory cell infiltration^64^. These effects are likely generalizable to other pathogens that cause pneumonia as influenza A virus (IAV) lacks tropism for cells in the skeletal muscle because the sialic acid moieties that serve as receptors for the virus are expressed only in airway epithelial cells, IAV^43,65,66^. Furthermore, in critically ill patients the severity of muscle atrophy correlates with serum levels of CRP, a transcriptional target of IL-6 in humans, and pharmacologic inhibition or genetic deletion of IL-6 prevents skeletal muscle loss after IAV infection in mice^67^.

We found that *Cx3cr1* and *Mertk* are necessary to localize a population of tissue-resident skeletal muscle macrophages with muscle fibers, satellite cells, FAPs, and endothelial cells, and the loss of this niche with aging is prevented by functional overexpression of MERTK. These findings align with findings from others who have causally linked spatially-localized signaling pathways to the loss of satellite cell proliferation after direct muscle injury in old mice. For example, signaling by NOTCH ligand requires direct interactions between cells in *trans*^10,11,68^. Furthermore, signaling by FGFs^17,18^, TGF-β and Wnts^19,20^, and activators of p38 MAPK pathways^21–24^ is typically restricted to juxtracrine or paracrine interactions^58^. Surprisingly, we did not observe expression of any of these ligands except TGF-β in skeletal muscle macrophages. Instead, these ligands were primarily expressed by FAPs and endothelial cells. These results could be explained if macrophages remove an inhibitory signal in the niche that precludes satellite cell proliferation. Support for this hypothesis comes from our own studies in cultured cells and the finding that microglia require *Mertk* and *Cx3cr1* for synaptic pruning during development, and from zebrafish models of skeletal muscle injury where monocyte-derived skeletal muscle macrophages are required to remove membrane patches on myofibers before satellite cell proliferation.

Heterochronic parabiosis has been shown to reverse the impaired satellite cell proliferation after direct tissue injury in old mice^11,55^. In contrast, heterochronic parabiosis did not reverse the age-related loss of myofiber-associated macrophages with aging. These differences might be explained by macrophage ontogeny. We used genetic lineage tracing to show monocyte-derived macrophages are not recruited to the skeletal muscle after injury^44^ and our findings suggest MERTK expression on tissue-resident skeletal muscle macrophages is required for satellite cell proliferation after IAV infection. In contrast, after direct muscle injury by toxins or pathologic genetic mutations, the release of damage-associated molecular patterns recruits monocytes from the circulation to the skeletal muscle where they differentiate into monocyte-derived skeletal muscle macrophages that are necessary for repair^37,39,69^. MERTK might also be important in monocyte-derived skeletal muscle macrophages. In a murine model of myocardial infarction, a loss or gain of function in Mertk in monocyte-derived macrophages was sufficient and necessary to worsen or ameliorate infarct size^47^. In old parabionts of a heterochronic pair, ∼40% of these circulating monocytes originate from the circulation of the young parabiont.

Our study has important limitations. We found that old *Mertk(CR)* mice were resistant to age-related frailty. Interestingly, however, age-related sarcopenia as measured by cross-sectional area, grip strength and voluntary wheel running was similar in old *Mertk(CR)* and old wild type mice. These findings are consistent with previous reports suggesting age-related sarcopenia is independent of satellite cells^70,71^. As MERTK is ubiquitously expressed in tissue-resident macrophages, further study of age-related phenotypes in these mice is warranted. Also, our mice voluntarily ran on wheels 2 weeks before and after IAV infection, suggesting this moderate level of exercise is not sufficient to allow recovery of skeletal muscle function after influenza A pneumonia in old mice^72^. However, we cannot exclude a role for long-term exercise during normal aging on the localization of skeletal muscle macrophages or skeletal muscle recovery from IAV-induced sarcopenia. Finally, our study is limited to mice, further studies using emerging spatial molecular approaches should focus on age-related changes in skeletal muscle macrophage localization in muscle biopsy samples from older individuals.

### Resource availability

#### Lead contact

Further information and requests for resources and reagents should be directed to and will be fulfilled by the lead contact, G.R. Scott Budinger

#### Materials availability

All unique reagents generated in this study are available from the lead contact.

## Supporting information

Movie S1 Young WT

Movie S2 Old WT

Movie S4 Old Mertk(CR)

Xenium Probes List

## Acknowledgments

We thank the following core facilities at Northwestern University: Pulmonary NextGen Sequencing Core, Northwestern University Behavioral Phenotyping Core, Northwestern University Robert H. Lurie Cancer Center RHLCCC Flow Cytometry Facility supported by NCI CA060553, Mouse Histology and Phenotyping Laboratory, and Northwestern University Center for Advanced Microscopy, supported by NCI CCSG P30-CA060553 awarded to the RHLCC. Some elements in Figure 1, 2 3 and supplemental 2 and 4 were generated with <u>BioRender.com</u>. This work was supported by the National Institutes for Health (NIH) grants P01AG049665 to J.I.S., N.S.C., A.V.M., and G.R.S.B.; R01HL173987 to J.I.S., N.S.C., and L.A.D.; 1F31AG071225 to R.A.G.; P01HL15499803 to N.S.C.; R01HL122309 to E.B.T., U19AI135964, P01HL154998, R01HL158139, R21AG075423 to A.V.M and G.R.S.B; U19AI181102, R01HL153312, R01ES034350 to A.V.M; U54AG079754, R01HL147575, R01HL158139, R01HL147290, R21AG075423 and U19AI135964 to G.R.S.B. R.A.G. was additionally supported by the Simpson Querrey Fellowship in Data Science as the Kimberly Querrey Fellow, and Schmidt Science Fellows program, in partnership with the Rhodes Trust, and G.R.S.B. by a Chicago Biomedical Consortium grant, Northwestern University Dixon Translational Science Award, Simpson Querrey Lung Institute for Translational Science, and Veterans Administration award no. I01CX001777.

## Author contributions

C.E.R., A.V.M and G.S.R.B. conceptualized the study with J.I.S, and interpreted the data and wrote the manuscript with the input of co-authors and assistance from N.S.C and L.A.D. C.E.R. performed experiments with the assistance of L.C.W., Z.L., F.C., R.A.N., R.A.G., and R. P.A. H.A-V. assisted in single-cell sequencing. L.L and K.J.S. analyzed single-cell RNA-seq data. E.G.B and W.T.P. led the mouse work and developed *Cx3cr1^ER-Cre/+^;zsGreen;Mertk^fl/fl^*and *Cx3cr1^Cre/+^;Mertk^fl/fl^* mouse lines, E.B.T. provided Mertk(CR) mice. C.W. performed behavioral phenotyping experiments and provided assistance interpreting results and editing the manuscript. A.V.M., M.J. S., and R.A.N. performed processing and analysis of Xenium spatial transcriptomics data.

## CONTACT FOR REAGENT AND RESOURCE SHARING

Please contact Dr. Budinger or Dr. Misharin for reagent or resource sharing

## EPERIMENTAL MODEL AND SUBJECT DETAILS

### Mouse Models

All mouse procedures were approved by the Institutional Animal Care and Use Committee at Northwestern University (Chicago, IL, USA). All strains including wild-type mice are bred and housed at a barrier- and specific pathogen–free facility at the Center for Comparative Medicine at Northwestern University. All experiments were performed with littermate controls. Number of animals per group was determined based on our previous publications. Mice were housed at the Center for Comparative Medicine at Northwestern University, in microisolator cages, with standard 12 hr light/darkness cycle, ambient temperature 23°C and were provided standard rodent diet (Envigo/Teklad LM-485) and water *ad libitum*. Four and eighteen months old C57BL/6J mice were obtained from the NIA colony. The C57BL/6J, *Cx3cr1^ER-Cre^* mice^73^ and ZsGreen reporter^74^ mice were obtained from Jackson laboratories (Jax stocks 000664, 020940 and 007906, correspondingly) as were Cx3cr1*^gfp/gfp^* mice (Jax stock 005582). Mertk(CR) mice were provided by Dr. Ira Tabas PMID 27199481. Mertkfl/fl mice were provided by Dr. Carla Rothlin PMID 28495875. Mertk-/- mice were obtained from the Jackson Laboratory (Strain #:038128).

## METHODS DETAIL

### Influenza-A infection

Murine-adapted, influenza A virus (A/WSN/33 [H1N1]) was provided by Robert Lamb, Ph.D., Sc.D., Northwestern University, Evanston, IL. Mice were anesthetized with isoflurane, their lungs were intubated orally with a 20-gauge Angiocath (Franklin Lanes, NJ) through which we administered either sterile, phosphate-buffered saline (S) or mouse-adapted influenza A virus (IAV) (50 μL) as previously described^66^. Various doses in a range of 10-100 pfu/mouse were tested to identify a dose that caused 20-40% mortality in young adult or old mice (Figure S1). We continuously observed mice infected with influenza A virus for signs of distress (slowed respiration, failure to respond to cage tapping, failure of grooming, huddling and fur ruffling). Mice that developed these terminal symptoms were sacrificed and the death was recorded as an influenza A-induced mortality. Weight was measured prior to influenza infection and prior to harvest.

### Measurement of muscle dysfunction

Forelimb skeletal muscle strength was assessed using a digital grip strength meter (Columbus Instruments, Columbus, OH) before harvest as described^66^. Grip strength was measured in each animal six successive times, and the average value for each mouse was recorded. The same operator performed all tests. After terminal anesthesia with Euthasol (pentobarbital sodium/phenytoin sodium), the soleus and *extensor digitorum longus* (EDL) muscles were excised and tendon was trimmed to a standard level under a microscope to assure precise weight measurements. The muscles were then blotted dry and weighed. Muscles were either frozen in liquid nitrogen-cooled isopentane for cryosectioning, snap-frozen in liquid nitrogen for protein extraction, or minced for flow cytometric analysis.

### Voluntary Wheel Running

Mice, housed 2-3 per cage, were provided with a trackable, low-profile saucer wheel (Med Associates, Inc., St. Albans, Vermont) 24 hours per day for 14-28 days during which time they developed consistent running patterns. After this equilibration period, all mice were infected with IAV or PBS at an age-appropriate dose (Figure S1C). Wheel rotations were gathered in one-minute increments via a wireless hub, and analyzed using wheel manager software (Med Associates, Inc., St. Albans, Vermont). Distances travelled per mouse per 24 hours (8 pm-8 pm) were calculated for each night. Data were collected as the average of 3 mice per cage for 6 cages each, and daily running distance is expressed relative to the average nightly distance in the averaged −5 days before infection.

### Frailty Assessment

The Frailty Index (FI) was scored as described previously^75^. This index includes 31 health-related deficits were assessed for each mouse. Measures across the integument, physical/musculoskeletal, oscular/nasal, digestive/urogenital, and respiratory systems were scored as 0, 0.5 and 1 based on the severity of the deficit. Total score across the items was divided by the number of items measured to give a frailty index score between 0 and 1.

### Behavioral Phenotyping

All assessments were performed by the Behavioral Phenotyping Core at Northwestern University.

Open Field Test: The dimension of the open field (OF) chamber was 60 cm × 60 cm × 30 cm. Animals were placed in the middle of the arena at the start of the test. They remained in the arena for 10 minutes and their positions were tracked using LimeLight software (Actimetrics). The center region was defined as an area that covered 44.4% of total arena (artificial dimension: 4×4 inside vs. 6×6 total arena).

Elevated zero-maze (EZM) test: The EZM consisted of a gray annular runway (5.5 cm width, 56 cm outer diameter, and 33 cm above ground level) inside a soundproof testing chamber. Two opposing 90° quadrants of the track were surrounded by inner and outer walls of gray polyvinyl-chloride (14 cm high). At the start of the test animals were placed at one end of the closed runway. Animals were allowed to stay in the maze for 10 minutes and their positions were tracked using LimeLight software.

DigiGait analysis: DigiGait imaging system (Mouse Specifics, Inc, Boston) was used for the evaluation of age-related changes in gait. Gait dynamics were recorded and analyzed, using Digigait Imager 16 software. Briefly, to acclimatize animals to the testing environment, the mouse was introduced to treadmill before the motorized belt began moving. Mice walked on the treadmill belt and recordings were taken at three gradually increasing speeds, 10 cm/s, 17 cm/s, and 24 cm/s. A 3-minute interval between recordings was set up to ensure mice were not overexerted. The gait dynamics were collected on video taken from a high-speed digital video camera mounted underneath the rig. For dynamic gait analysis, the video was reviewed and the noise was manually corrected from dynamic gait signals. A minimum of 4 seconds (150 frames/s) of the movie was required. Each analysis was performed twice for each mouse and speed.

### Immunohistochemistry for Fiber Size Assessment and Macrophage Localization

Tibialis anterior (TA) soleus and extensor digitorum longus (EDL) serial transverse cryosections (8 μm) were mounted on glass slides. Sections were fixed in ice-cold acetone, permeabilized, and blocked. Immunostaining was performed using primary antibodies against the following epitopes laminin (1:50 dilution; Sigma; Catalog: L9393) or CD68 (1:400 dilution; Invitrogen Catalog PA5-78896) or (1:200 dilution; Invitrogen Catalog 14-0681-82, or Peg3 (1:200 dilution; Invitrogen PA5-99683) followed by Alexa Fluor-conjugated secondary antibodies (1:300 dilution; Life Technologies; Catalog: A-11011). Images were acquired with a Zeiss LSM 510 confocal microscope using a 20x objective (Northwestern University Center for Advanced Microscopy) and analyzed using Zeiss LSM5 Image Browser software. Fiber size was studied by measuring the fibers’ minimal inner diameter (at least 100 fibers per muscle), defined as the minimum diameter from inner border to inner border, passing through the center of the muscle fiber. This parameter has been shown to be very insensitive to deviations from the “optimal” cross-sectioning profile, as compared with direct measure of fiber cross-sectional area^66^. Cross-sectional area (CSA) was calculated using this diameter, and results were expressed as mean CSA ± S.E. Alternatively, soleus and TA muscles were sectioned and stained for immunohistochemical detection of CD68 in the Northwestern University Mouse Histology and Phenotyping Laboratory.

### Flow Cytometry and Cell Sorting

Mice were fully-perfused through the right ventricle with 20 ml of PBS, and hind-limb muscle was removed. The tissue was cut into small pieces with scissors, transferred into C-tubes (Miltenyi, Auburn, CA) and processed with a Skeletal Muscle Dissociation kit according to manufactures instructions (Miltenyi, Auburn CA). To achieve a single-cell suspension, 2 rounds of dissociation were performed using a GentleMACS dissociator (Miltenyi), followed by 3 rounds of filtration through Falcon 100-μm, 70-μm then 40-μm nylon mesh filter units (Thermo Fisher #352340, 352350 and 352360 respectively) into polypropylene 50ml Falcon tubes, followed by centrifugation at 1300 rpm in an Eppendorf 5810R centrifuge for 15 minutes. Any remaining red blood cells were lysed by resuspending the pellet in 1ml/tube BD Pharm Lyse (BD Biosciences, San Jose, CA), and were transferred to flow tubes, pelleted by low-speed centrifugation 5 minutes, and resuspended in 0.5ml/tube HBSS containing 0.5 μl/tube viability dye eFluor 506 (eBioscience, San Diego, CA). Cells were stained in viability dye for 15 minutes in the dark at room temperature, followed by 2X washes in 1ml MACS buffer (Miltenyi, Auburn, CA). After live/dead cell staining, pellets are resuspended in 150 μl/tube FcBlock Anti-mouse CD16/CD32 (Invitrogen) for 30 minutes in the dark at 4°C to block non-specific binding. After blocking, cells were divided to two separate tubes at 50 μl each, then stained with a mixture of fluorochrome-conjugated antibodies in 50 μl/tube and incubated for an additional 30 minutes at 4°C in the dark. After antibody incubation, the cells were washed 2X in MACS buffer, and then fixed in 2% paraformaldehyde for 15 minutes at room temperature in the dark, washed 2X in HBSS, resuspended in 200 μl HBSS, then stored at 4°C overnight. Data were acquired on a BD FACSymphony A-5 laser analylzer using BD FACSDiva software, and compensation and data analyses were performed using FlowJo software (TreeStar, Ashland, OR). Prior to cytometric analysis, 50 μl 123count eBeads (Invitrogen) were added to 200 μl each sample to allow accurate cell counts/gm of tissue. “Fluorescence minus one” controls were used when necessary. Cell populations were identified using sequential gating strategy (See Figure S2B, C). Cell sorting was performed at Northwestern University RLHCCC Flow Cytometry core facility on SORP FACSAria III instrument (BD) with the same optical configuration as LSR II, using a 100-µm nozzle and 40 psi pressure.

### BMDM isolation and co-culture experiments

Bone marrow was isolated from mice and plated in 10 cm tissue culture dishes (Corning 353003). To induce differentiation into macrophages, cells were cultured in DMEM (Corning 10-013-CV), 10% fetal bovine serum (Corning 35-001-CV), 1% antibiotic/mycotic (ThermoFisher, catalog no. 15-240-062) 1% HEPES (ThermoFisher, catalog no. MT25060CI). supplemented with 20ng/ml M-CSF (Peprotech, catalog no.315-02) at 37 °C with 5% CO2. The medium was changed every 3 days, and BMDMs were harvested by trypsin-EDTA, counted and plated in equal numbers (0.25 X 10^5^ cells/ml) with C2C12 myoblasts (ATCC, Manassas, VA: CRL-1772) for co-culture experiments in triplicate wells of 6-well plates (Corning 353224) containing sterilized square glass cover slips (VWR 48386 067). Triplicate wells of C2C12 cells (0.25 X 10^5^ cells/ ml) but without added BMDM served as controls. Two hours post co-plating, after cells had attached, three coverslips per condition were removed and assessed as day 0 for C2C12 plated alone, or in combination with BMDM. Cells were fixed in 4% paraformadehyde for 15 minutes, followed by permeabilization, blocking incubation in primary antibodies F4/80-Biotin (1:200 Biolegend catalog 123106)) and MYH3 (1:400 dilution; Santa Cruz catalog number sc-53091) Overnight at 4°C in a humidified chamber. Secondary antibody incubation was performed at room temperature with streptavidin-AF594 (1:500 ThermoFisher catalog S11227) followed by nuclear staining with Hoechst (1:10,000 Invitrogen catalog 33342). Five representative fields were imaged for each coverslip and counted for total nuclei (all Hoechst positive) or BMDM nuclei (F4/80 positive) or fused C2C12 nuclei (nuclei fused two or more expressed as a percentage of total nuclei minus BMDM). The five fields for each measure were averaged to represent single data points for each coverslip. Similar assessments were performed at day 2, 4 and 7.

### Western Blot Analysis

Quadriceps muscle tissue was homogenized with a Tissue Tearor (BioSpec Products, Inc) for 1 minute in ice-cold lysis buffer (Nonidet P-40 1%, glycerol 10%, NaCl 137 mM, Tris-HCl pH 7.5 20 mM) containing protease (1 Complete Mini, EDTA-free tablet, Roche) and phosphatase (sodium fluoride 30 mM, β-glycerophospate 250 mM, sodium orthovanadate 1mM) inhibitors. Samples were centrifuged at 15,000 rpm for 10 minutes at 4°C and the supernatant was collected. Protein concentrations were determined with Protein Assay Dye (Bio-Rad) in a 96 well plate measured against BSA protein standard dilutions at 595 nm with a Bio-Rad iMark microplate reader (Bio-Rad, Hercules CA). Equal amount of protein (40-80 total μg) were loaded on a 9% SDS-PAGE gel, and Western blot analysis was performed as described previously^66^. Incubation with primary antibodies was performed overnight at 4°C. Immunoblots were quantified by densitometry using Image J 1.46r (National Institutes of Health, Bethesda, MD) or Image Studio Software (LI-COR Inc., Lincoln, NE). The following antibodies were used: Rabbit Monoclonal to Fbx32 (Abcam; catalog: ab168372; 1:1000) or Rabbit monoclonal to GAPDH (D16H11) (Cell Signaling; catalog: 5174s; 1:1000).

### scRNA-seq Library Preparation and Generation

Harvest of muscle tissue was performed with two mice each from each group. Three harvests were done over three consecutive days: 9, 10 or 11 dpi. Mice were fully perfused through the right ventricle with 20 ml of PBS, and hind limb muscle was removed. Tissue was cut into small pieces with scissors, transferred into C-tubes (Miltenyi), and processed with a Skeletal Muscle Dissociation Kit according to manufacturer’s instructions (Miltenyi). To achieve a single-cell suspension, two rounds of dissociation were performed using a GentleMACS dissociator (Miltenyi), followed by filtration through Falcon 70-μm, and then 40-μm nylon mesh filter units (Thermo Fisher #352340, 352350, and 352360, respectively) into polypropylene 50-ml Falcon tubes, followed by centrifugation at 300 rcf in an Eppendorf 5810R centrifuge for 15 min. Any remaining red blood cells were lysed by resuspending the pellet in 1ml/tube BD Pharm Lyse (BD Biosciences), and were transferred to flow tubes, pelleted by low-speed centrifugation 5 min, resuspended and combined for two mice per condition into 100 μl/tube FcBlock Anti-mouse CD16/CD32 (Invitrogen) for 30 min in the dark at 4°C to block non-specific binding. After blocking, cells were stained in 100 μl each in a mixture of fluorochrome-conjugated antibodies. After incubation at 4 °C for 30 min, cells were washed with 2 ml MACS buffer, pelleted by centrifugation, and resuspended in 500 μl MACS buffer with 2 μl SYTOX Green viability dye (ThermoFisher). Cell sorting was performed at Northwestern University RLHCCC Flow Cytometry core facility on SORP FACSAria III instrument (BD) using a 70-μm nozzle at 20 psi. 10,000 cells each of monocyte/macrophage (MO/MP), fibroadipogenic progenitor (FAP), or satellite cells (MuSC) were sorted together into 300 μl of 2% BSA in PBS for single-cell RNA-seq. Myeloid cell populations cell populations were defined as CD45+, CD11b+, Ly6G-, SiglecF-, and assessed for CD64 and MHCII as additional macrophage markers. FAP were defined as CD45-, CD31-, a7 integrin- and Sca-1+, and MuSC were defined as CD45-, CD31-, a7 integrin+ and Sca-1-. Following sorting, cells were counted and assed for viability. Cells were washed, and each tube containing MO/MP, FAP and MuSC from each of the 4 groups was resuspended in 100ul of cell multiplexing oligonucleotides (CMO) according to the kit protocol from 10X Genomics. Libraries were generated using 10x genomics 3’ v3 chemistry.

### scRNA-seq Analysis

Initial data processing was performed using the Cell Ranger version 6.1.2 pipeline (10x Genomics). Post-processing, including filtering by number of genes expressed per cell, was performed using the Seurat package V4.3.0.1 and R 4.3.2 (62, 63). Briefly, gene counts from sorted FAPs, muscle macrophages, and muscle satellite cells were demultiplexed using cell multiplexing oligonucleotide (CMO) hashtag antibodies with the “CLR” normalization method and the HTODemux function in Seurat. Counts were filtered for cells expressing more than 200 genes and genes expressed in more than 3 cells. After aggregation in Seurat, doublet clusters, clusters dominated by a single biosample, or low-quality clusters expressing primarily mitochondrial genes were removed prior to integration. Raw gene counts were normalized using SCTransform and integrated across biosample using scVI with 2000 highly variable genes. Following clustering with the Leiden algorithm and visualization with UMAP, initial clusters were subjected to inspection and merging based on similarity of marker genes and a function for measuring phylogenetic identity using BuildClusterTree in Seurat. Identification of cell clusters was performed on the final object guided by marker genes. Differential gene expression analysis was performed for each cluster between cells from mice of each condition using DESeq2. Heatmaps, UMAPs, violin plots, dot plots, and boxplots were generated using Seurat.

Differential cell type abundance analysis was performed within the 3 main sorted cell populations of muscle satellite cells, FAPs, and muscle macrophages. Monocytes were excluded from differential abundance analysis within muscle macrophages. Statistical comparisons were performed using the Wilcoxon rank sum test with adjustment for multiple comparisons (FDR < 0.05).

### Spatial Transcriptomics

Cell segmentation was performed with 10X Xenium onboard cell segmentation (Xenium Ranger v1.7.6.0, 10x Genomics). DAPI stained nuclei from the DAPI morphology image were segmented and boundaries were consolidated to form non-overlapping objects. To approximate cell segmentation, nuclear boundaries were expanded by 15 μm or until they reached another cell boundary. For each sample, Xenium generated an output file of transcript information including x and y coordinates, corresponding gene target, assigned cell, a binary flag for whether the transcript was expressed over a nucleus, and quality score. Low-quality transcripts (qv < 20) and transcripts corresponding to blank/negative probes were removed. Seurat v5 was used to perform further quality filtering and visualization on nuclei gene expression data. Nucleus by gene count matrices were created for each sample based on expression of transcripts that fell within segmented nuclei. A single merged Seurat object was created for all samples based on these count matrices and metadata files with nuclei coordinates and area. Nuclei were retained according to the following criteria: ≥ 2 unique genes. Because Xenium outputs coordinates based on each slide, which results in samples with shared coordinates across multiple slides, the nuclei coordinates were manually adjusted for visualization so that no samples overlapped during plotting. These adjusted nucleus coordinates were added to the Seurat object and h5ad object as meta-data columns. Seurat v5 was used to perform dimensionality reduction, clustering, and visualization. Gene expression was normalized per cell using Seurat’s scTransform function. Dimensionality reduction was performed with PCA. Nuclei were clustered based on this dimensionality reduction using the Louvain algorithm. Uniform Manifold Approximation and Projection (UMAP) plots of the data were generated using the same number of PCs as used for PCA. Cells were annotated using a combination of marker genes, generated using Seurat v5’s FindMarkers function, and spatial information including cell morphology and position. The CellCharter package in python was used to partition cells into 5, 8 and 12 spatial niches using k-means clustering based on the transcriptomic expression of the nearest 6 neighboring cells.

## QUANTIFICATION AND STATISTICAL ANALYSIS

### Statistical Analysis

Differences between groups were determined according to ANOVA. When ANOVA indicated a significant difference, individual differences were examined using *t* tests with a Tukey correction for multiple comparisons, as indicated. All analyses were performed using GraphPad Prism, version 5.00 (GraphPad Software, San Diego CA). Data are shown as means ± SEMs. *P* < 0.05 was considered statistically significant in all tests.

