## Supplemental Figures for "Tissue-resident skeletal muscle macrophages promote recovery from viral pneumonia-induced sarcopenia in normal aging"

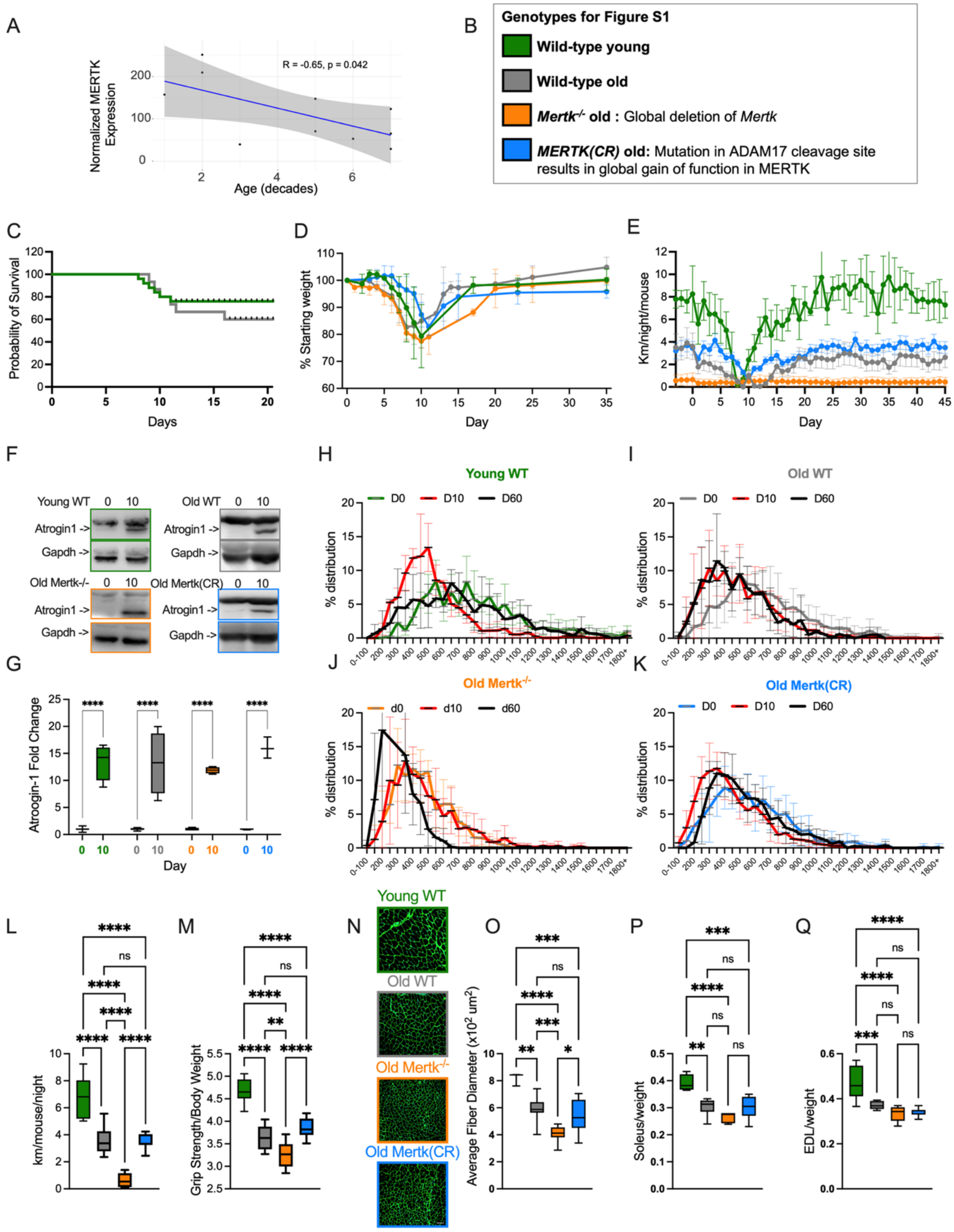


**Figure S1: *Mertk(CR)* mice are resistant to some age-related phenotypes.** See also Figure 1

(A) Pearson's correlation of DESeq2 normalized MERTK expression counts in macrophages from the Human Muscle Ageing Atlas vs age in decades on a per sample level.
(B) Murine strains and ages used in Figure S1
(C) Survival after intratracheal instillation of 20 pfu IAV/animal young wild-type mouse, or 5 pfu IAV/animal old wild-type mouse (n=16-20 mice per group).
(D) Weight as percentage of starting weight before influenza A virus (IAV) infection for young adult wild-type, old wild-type, old *Mertk*$^{-/-}$, and old *Mertk(CR)* mice (n=7-12 mice/group).
(E) Daily voluntary wheel running distance (km/day) before and after IAV infection for young adult wild-type, old wild-type, old *Mertk*$^{-/-}$, and old *Mertk(CR)* mice for young adult wild-type, old wild-type, old *Mertk*$^{-/-}$, and old *Mertk(CR)* mice (n=9-12 mice per group).
(F) Representative Western blot of ATROGIN-1 expression in the quadriceps muscle for young adult wild-type, old wild-type, old *Mertk*$^{-/-}$, and old *Mertk(CR)* mice. Arrows denote ATROGIN-1, or GAPDH loading control
(G) Box plots of ATROGIN-1 fold change in quadriceps muscle normalized to GAPDH for young adult wild-type, old wild-type, old *Mertk*$^{-/-}$, and old *Mertk(CR)* mice. (n=3-6) Two-way ANOVA was used to determine statistical significance.
(H) Histograms of fiber size distribution at day 0, day 10 or day 60 after influenza A infection (n=5 mice per group) for young adult wild-type mice (n=5-12 mice per group).
(I) Histograms of fiber size distribution at day 0, day 10 or day 60 after influenza A infection (n=5 mice per group) for old wild-type mice (n=5-12 mice per group).
(J) Histograms of fiber size distribution at day 0, day 10 or day 60 after influenza A infection (n=5 mice per group) for old *Mertk*$^{-/-}$ mice (n=5-12 mice per group).
(K) Histograms of fiber size distribution at day 0, day 10 or day 60 after influenza A infection (n=5 mice per group) for old *Mertk(CR)* mice (n=5-12 mice per group).
(L) Box plots depicting per mouse averaged nightly running wheels distances from 5 days for young adult wild-type, old wild-type, old *Mertk*$^{-/-}$, and old *Mertk(CR)* mice (8-10 wheels per group). Ordinary one-way ANOVA used to determine statistical significance.
(M) Box plots depicting averaged forearm grip strength corrected for weight of the mouse for young adult wild-type, old wild-type, old *Mertk*$^{-/-}$, and old *Mertk(CR)* mice (8-12 per group). Ordinary one-way ANOVA used to determine statistical significance.
(N) Representative images of laminin-stained tibialis anterior muscle transverse cross sections for young adult wild-type, old wild-type, old *Mertk*$^{-/-}$, and old *Mertk(CR)* mice (scale bar = 50mm),
(O) Box plot graphs depicting average cross sectional fiber area for young adult wild-type, old wild-type, old *Mertk*$^{-/-}$, and old *Mertk(CR)* mice (n=5-12 mice per group). Ordinary one-way ANOVA used to determine statistical significance.
(P) Box plots depicting average soleus muscle wet weight corrected for mouse total body weight for young adult wild-type, old wild-type, old *Mertk*$^{-/-}$, and old *Mertk(CR)* mice (n=5-9 mice per group). Ordinary one-way ANOVA used to determine statistical significance.
(Q) Box plots depicting average extensor digitorum longus muscle wet weight corrected for total body weight for young adult wild-type, old wild-type, old *Mertk*$^{-/-}$, and old *Mertk(CR)* mice (n=5-9 mice per group). Ordinary one-way ANOVA used to determine statistical significance

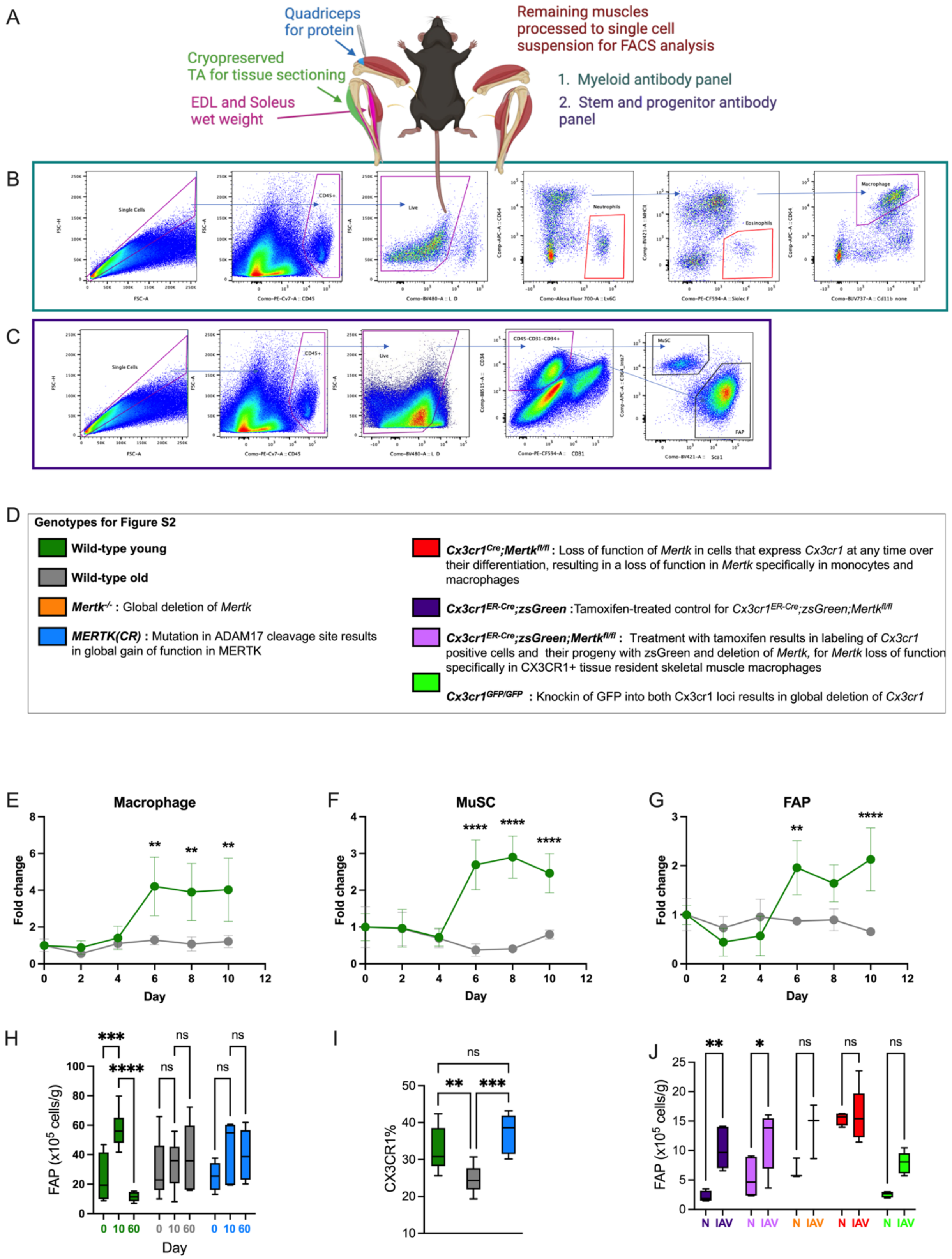
A
Quadriceps for protein
Cryopreserved TA for tissue sectioning
EDL and Soleus wet weight
Remaining muscles processed to single cell suspension for FACS analysis
1. Myeloid antibody panel
2. Stem and progenitor antibody panel
B
Single Cells
CD45+
Live
Neutrophils
Eosinophils
Macrophage
C
Single Cells
CD45+
Live
CD45-CD31-CD34+
MuSC
FAP
D
Genotypes for Figure S2
Wild-type young
Wild-type old
Mertk-/- : Global deletion of Mertk
MERTK(CR) : Mutation in ADAM17 cleavage site results in global gain of function in MERTK
Cx3cr1Cre;Mertkfl/fl : Loss of function of Mertk in cells that express Cx3cr1 at any time over their differentiation, resulting in a loss of function in Mertk specifically in monocytes and macrophages
Cx3cr1ER-Cre;zsGreen :Tamoxifen-treated control for Cx3cr1ER-Cre;zsGreen;Mertkfl/fl
Cx3cr1ER-Cre;zsGreen;Mertkfl/fl : Treatment with tamoxifen results in labeling of Cx3cr1 positive cells and their progeny with zsGreen and deletion of Mertk, for Mertk loss of function specifically in CX3CR1+ tissue resident skeletal muscle macrophages
Cx3cr1GFP/GFP : Knockin of GFP into both Cx3cr1 loci results in global deletion of Cx3cr1
E
Macrophage
Fold change
Day
F
MuSC
Fold change
Day
G
FAP
Fold change
Day
H
FAP (x10^5 cells/g)
0 10 60
Day
I
CX3CR1%
J
FAP (x10^5 cells/g)
N IAV

**Figure S2: The expansion of macrophages and satellite cells during recovery from influenza A infection requires the expression of *Mertk* and *Cx3cr1* in tissue-resident skeletal muscle macrophages.** See also Figure 2B-E

(A) Schematic demonstrating isolation of muscle tissue for analysis for figures 1 and 2.
(B) Gating strategy for flow cytometry analysis of skeletal muscle macrophages.
(C) Gating strategy for muscle satellite cells (MuSC) and FAP.
(D) Murine genotypes used for these experiments.
(E) Single cell suspensions were generated from the muscles of young adult or old wild type mice and macrophages were quantified using flow cytometry 0, 2, 4, 6 8-, or 10-days post influenza A infection. Line graphs show fold change in macrophage number per gram of tissue compared to day 0. (n=3-6 mice per group). Two-way ANOVA was used to determine statistical significance.
(F) Single cell suspensions were generated from the muscles of young adult or old wild type mice and muscle satellite cells (MuSC) were quantified using flow cytometry 0, 2, 4, 6 8-, or 10-days post influenza A infection. Line graphs show fold change in macrophage number per gram of tissue compared to day 0. (n=3-6 mice per group). Two-way ANOVA was used to determine statistical significance.
(G) Single cell suspensions were generated from the muscles of young adult or old wild type mice and FAP cells were quantified using flow cytometry 0, 2, 4, 6 8-, or 10-days post influenza A infection. Line graphs show fold change in macrophage number per gram of tissue compared to day 0. (n=3-6 mice per group). Two-way ANOVA was used to determine statistical significance.
(H) FAP cells were quantified using flow cytometry from single cell suspensions of hindlimb muscles from young adult wild-type, old wild-type, and old *Mertk(CR)* mice at day 0 (naïve, N), or 10-day post IAV. Box plots show total numbers of FAP per gram of tissue (n=6-12 mice per group). Two-way ANOVA was used to determine statistical significance.
(I) Box plots show percentage of muscle macrophages that express CX3CR1 for young or old wild-type, or old *Mertk(CR)* mice. (n=6-8 mice per group). Ordinary one-way ANOVA used to determine statistical significance.
(J) FAP cells were quantified using flow cytometry from single cell suspensions of hindlimb muscles from $Cx3cr1^{ER\text{-}Cre/+}$*;zsGreen*, $Cx3cr1^{ER\text{-}Cre/+}$*;zsGreen;*$Mertk^{fl/fl}$, $Mertk^{-/-}$, $Cx3cr1^{Cre/+}$;$Mertk^{fl/fl}$ and $Cx3cr1^{GFP/GFP}$ mice at day 0 (naïve, N), or 10-day post IAV. Box plots show total numbers of FAP per gram of tissue (n=6-12 mice per group). Two-way ANOVA was used to determine statistical significance.

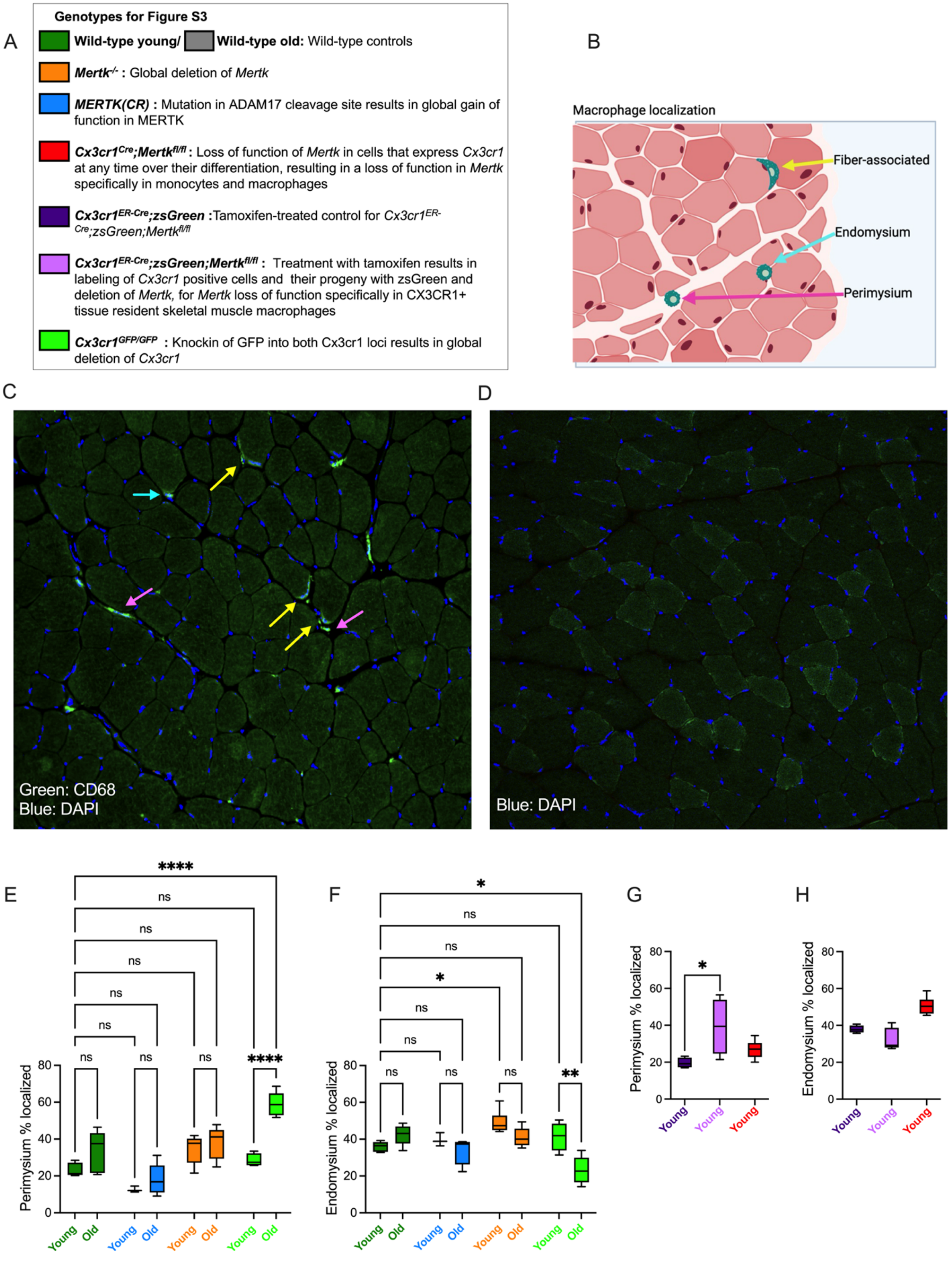
A
Genotypes for Figure S3
Wild-type young/ Wild-type old: Wild-type controls
Mertk-/- : Global deletion of Mertk
MERTK(CR) : Mutation in ADAM17 cleavage site results in global gain of function in MERTK
Cx3cr1Cre;Mertkfl/fl : Loss of function of Mertk in cells that express Cx3cr1 at any time over their differentiation, resulting in a loss of function in Mertk specifically in monocytes and macrophages
Cx3cr1ER-Cre;zsGreen :Tamoxifen-treated control for Cx3cr1ER-Cre;zsGreen;Mertkfl/fl
Cx3cr1ER-Cre;zsGreen;Mertkfl/fl : Treatment with tamoxifen results in labeling of Cx3cr1 positive cells and their progeny with zsGreen and deletion of Mertk, for Mertk loss of function specifically in CX3CR1+ tissue resident skeletal muscle macrophages
Cx3cr1GFP/GFP : Knockin of GFP into both Cx3cr1 loci results in global deletion of Cx3cr1
B
Macrophage localization
Fiber-associated
Endomysium
Perimysium
C
Green: CD68
Blue: DAPI
D
Blue: DAPI
E
Perimysium % localized
****
ns
ns
ns
ns
ns
ns
ns
ns
****
Young
Old
F
Endomysium % localized
*
ns
ns
*
ns
ns
ns
ns
ns
**
Young
Old
G
Perimysium % localized
*
Young
H
Endomysium % localized
Young

**Figure S3: The localization and expansion of tissue resident *Cx3cr1*+ macrophages and satellite cells after influenza infection requires *Cx3cr1* and *Mertk* in young mice, is lost in old mice, and is maintained in old *Mertk(CR)* mice.** See also Figure 2F-I.

(A) Murine genotypes used for these experiments.
(B) Schematic demonstrating localization of tissue resident macrophages within skeletal muscle cross sections**.**
(C) Tibialis anterior muscle cross section from young adult mice showing representative localizations muscle macrophages. Yellow arrows indicate fiber associated macrophages, pink arrows indicate macrophages in the perimysium and aqua labels indicate macrophages in the endomysium. All macrophages are indicated by CD68. For assessment of macrophage localization, 5-7, 10-micron sections were made per muscle, each section was imaged 3-5 times to cover the entire section, and every macrophage in each image (excluding epimysium because of its variable thickness and being inconsistently intact) was cumulatively scored for location with an average total of 150-200 macrophages per muscle. For each mouse/muscle the total counts by localization are expressed as a percent of total macrophages counted.
(D) Representative secondary antibody negative control for CD68.
(E) Box plots display the percent of total macrophages that are localized with the perimysium of naïve young adult mice or old mice from wild-type, *Mertk*$^{-/-}$ and *Cx3cr1*$^{GFP/GFP}$ . (n=3-5 mice per group) Two-way ANOVA was used to determine statistical significance.
(F) Box plots display the percent of total macrophages that are localized with the endomysium of naïve young adult mice or old mice from wild-type, *Mertk*$^{-/-}$ and *Cx3cr1*$^{GFP/GFP}$. (n=3-5 mice per group) Two-way ANOVA was used to determine statistical significance.
(G) Box plots display the percent of total macrophages that are localized with the perimysium in young adult *Cx3cr1*$^{ER\text{-}Cre;}$*zsGreen* and *Cx3cr1*$^{ER\text{-}Cre;}$*zsGreen; Mertk*$^{fl/fl}$ mice after treatment with tamoxifen by oral gavage 10 and 9 days before harvest, and in young adult *Cx3cr1*$^{Cre}$*; Mertk*$^{fl/fl}$ mice. (n=4-5 mice per group) Unpaired t test was used to determine statistical significance.
(H) Box plots display the percent of total macrophages that are localized with the endomysium in young adult *Cx3cr1*$^{ER\text{-}Cre;}$*zsGreen* and *Cx3cr1*$^{ER\text{-}Cre;}$*zsGreen; Mertk*$^{fl/fl}$ mice after treatment with tamoxifen by oral gavage 10 and 9 days before harvest, and in young adult *Cx3cr1*$^{Cre}$*; Mertk*$^{fl/fl}$ mice. (n=4-5 mice per group) Unpaired t test was used to determine statistical significance.

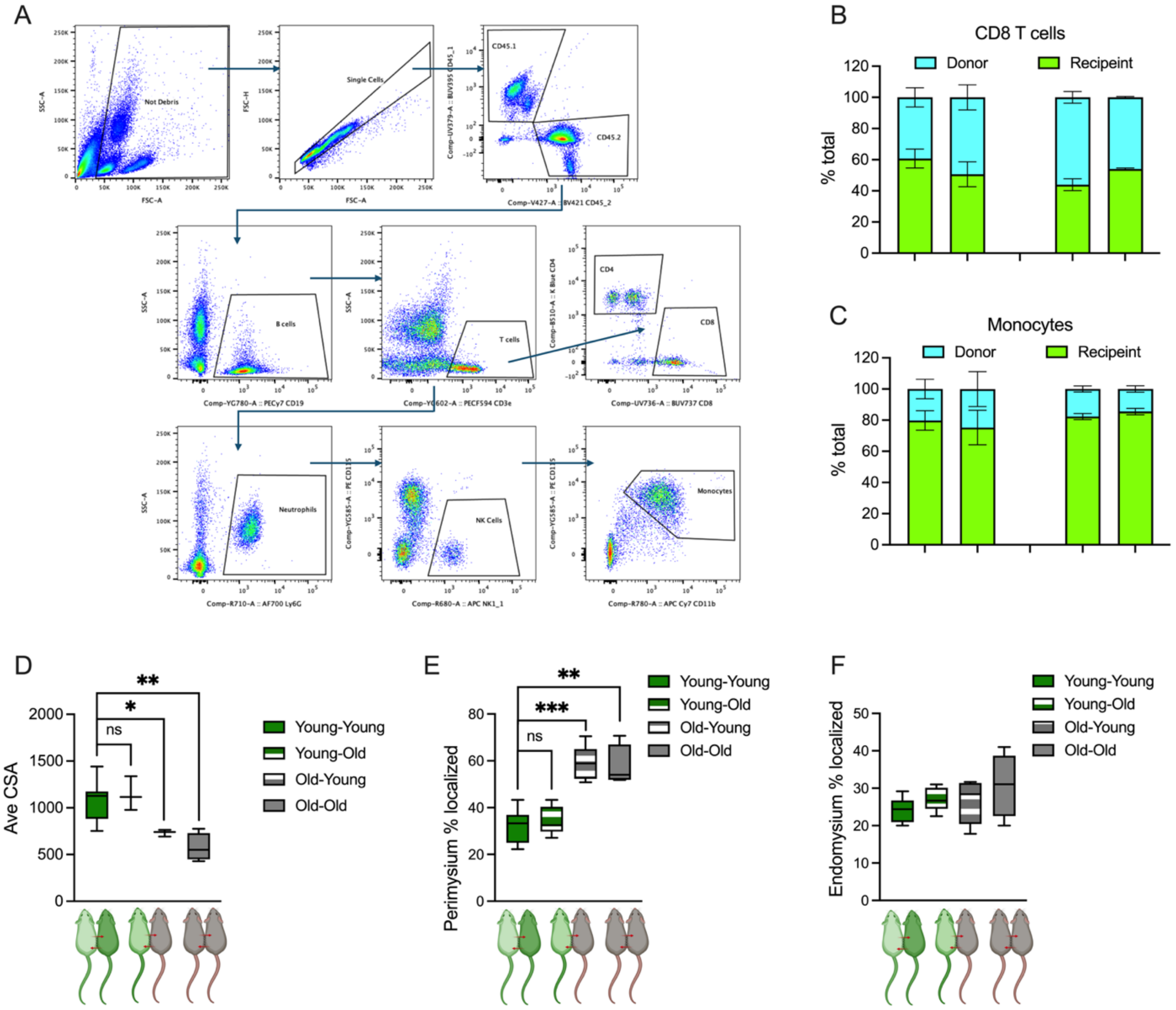


**Figure S4: Heterochronic parabiosis do not show alteration of tissue-resident muscle macrophage localization**. See also Figure 2J

(A) Representative gating for flow cytometry of immune cells from blood to assess chimerism in CD45.1-CD45.2 parabionts.

(B) Percent of donor-derived T cells in blood within parabionts for young CD45.1 (light green) parabiotic pairs with young CD45.2 (dark green) or old CD45.2 (grey) mice.

(C) Percent of donor-derived monocytes in blood within parabionts for young CD45.1 (light green) parabiotic pairs with young CD45.2 (dark green) or old CD45.2 (grey) mice.

(D) Young adult CD45.1 (light green) and CD45.2 young (dark green) or old (grey) mice were joined as heterochronic parabionts for 60 days as indicated. Box plot graphs depict average tibialis anterior muscle cross sectional fiber area (n=3-6 mice per group). Unpaired t test was used to determine statistical significance.

(E) Young adult CD45.1 (light green) and CD45.2 young (dark green) or old (grey) mice were joined as heterochronic parabionts for 60 days as indicated. Box plots display the percentage of total skeletal muscle macrophages localized in the perimysium (n=3-6 mice per group). Unpaired t test was used to determine statistical significance.

(F) Young adult CD45.1 (light green) and CD45.2 young (dark green) or old (grey) mice were joined as heterochronic parabionts for 60 days as indicated. Box plots display the percentage of total skeletal muscle macrophages localized in the endomysium (n=3-6 mice per group). Unpaired t test determined no statistical significance.

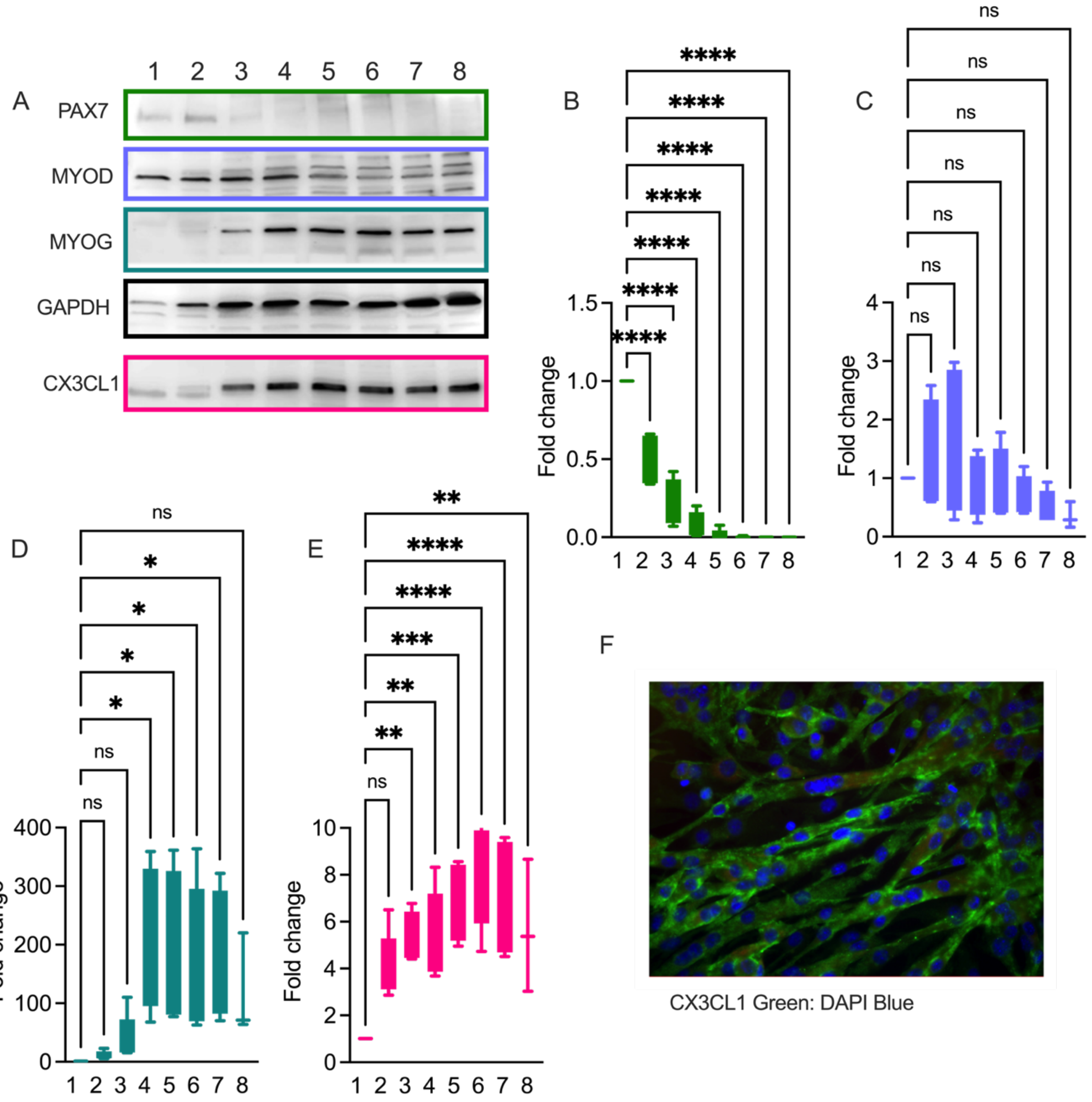


**Figure S5. CX3CL1 increases in C2C12 cells as they undergo proliferation and differentiation.** See also Figure 3

(A) Representative immunoblots showing expression of PAX7, MYOD, MYOG, GAPDH, and CX3CL1 in homogenates of C2C12 cells Day 1-8 of culture.
(B) Quantification of expression from immunoblots for PAX7 expression corrected for GAPDH expression (n=4) Ordinary one-way Anova used to determine significance compared to Day 1.
(C) Quantification of expression from immunoblots for MYOD expression corrected for GAPDH expression (n=4) Ordinary one-way Anova used to determine significance compared to Day 1.
(D) Quantification of expression from immunoblots for MYOG expression corrected for GAPDH expression (n=4) Ordinary one-way Anova used to determine significance compared to Day 1.
(E) Quantification of expression from immunoblots for CX3CL1 expression corrected for GAPDH expression (n=4) Ordinary one-way Anova used to determine significance compared to Day 1.
(F) Representative images of C2C12 myotubes at Day 6 of culture cells that had been previously treated stained with CX3CL1 (Green).

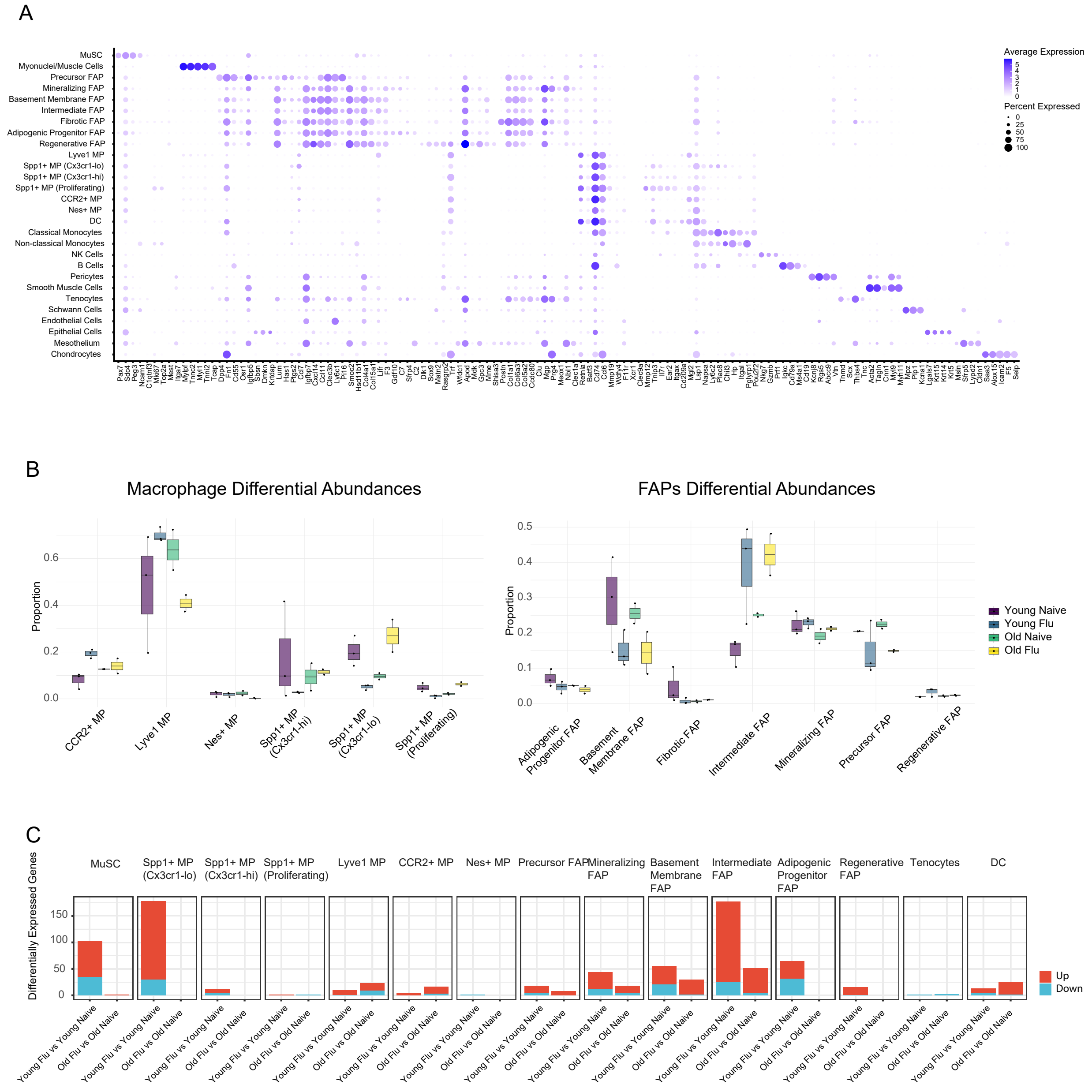


**Figure S6. Single cell RNA-sequencing of skeletal muscle from young adult and old mice before and 10 days after influenza A pneumonia.** Refers to Figure 4A

(A) Dot plot of genes used to support cell annotations.
(B) Comparison of the abundance of macrophages and FAP populations estimated from single cell RNA sequencing data of skeletal muscle homogenates between naive young adult and old mice and young adult and old mice 10 days after influenza A infection. There were no significant differences (FDR q<0.05).
(C) Pseudobulk differential gene expression analysis was used to measure the number of differentially expressed genes 10 days after influenza A infection in young adult and old mice across resolved cell populations (FDR q<0.05)

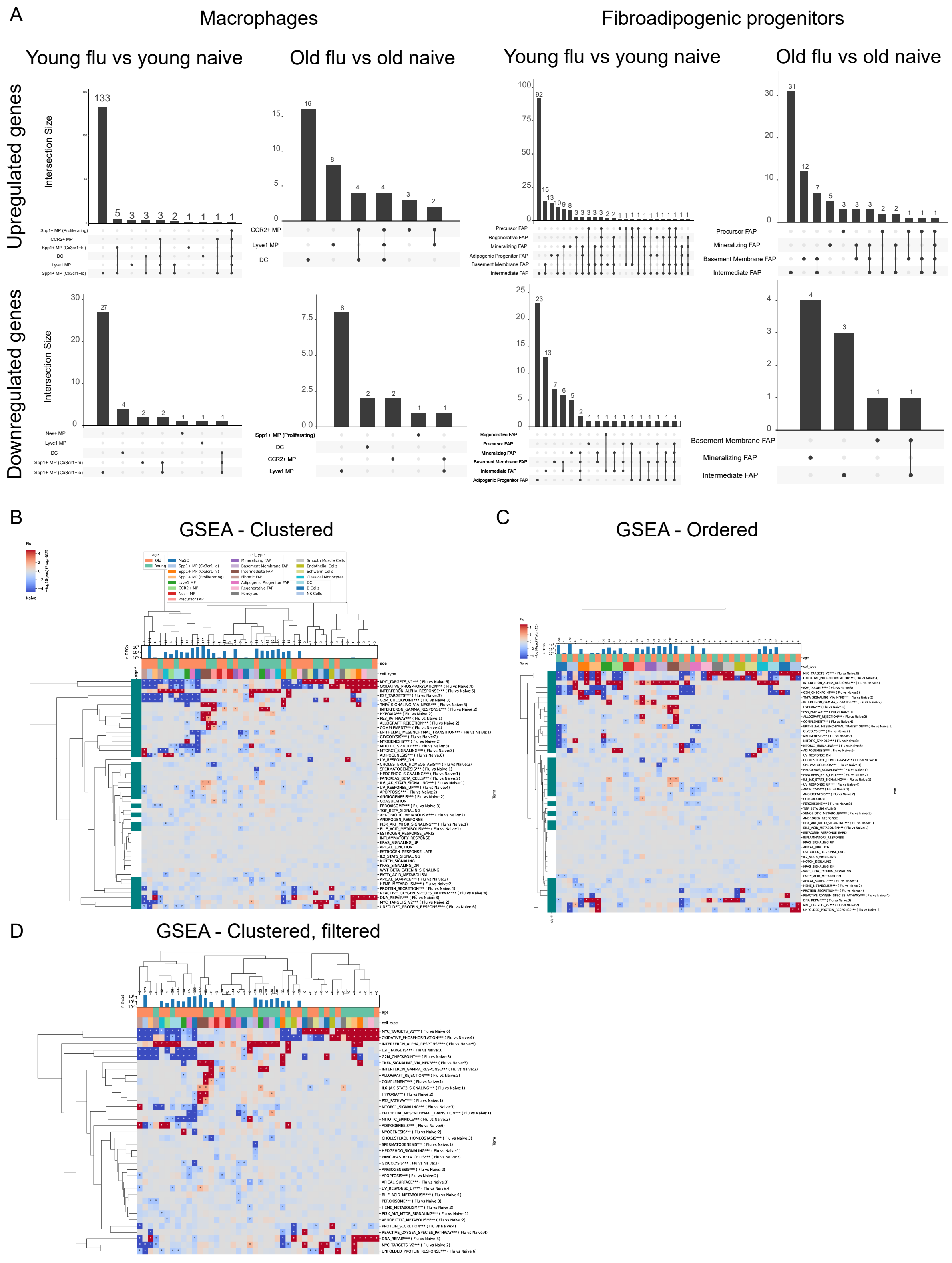


**Figure S7. Single cell RNA-sequencing of skeletal muscle from young adult and old mice before and 10 days after influenza A pneumonia.** Refers to Figure 4B.

(A) Upset plots showing overlap between differentially expressed genes in the absence of or 10 days after influenza A infection in young and old mice across resolved macrophage and FAP populations (FDR q<0.05).
(B) Heatmap showing GSEA where both rows (processes) and columns (cell types) are clustered.
(C) Heatmap showing GSEA where only rows (processes) are clustered.
(D) Heatmap showing GSEA processes that were significantly differentially expressed in the absence of or 10 days after influenza A infection in young adult and old mice where both rows (processes) and columns (cell types) are clustered.

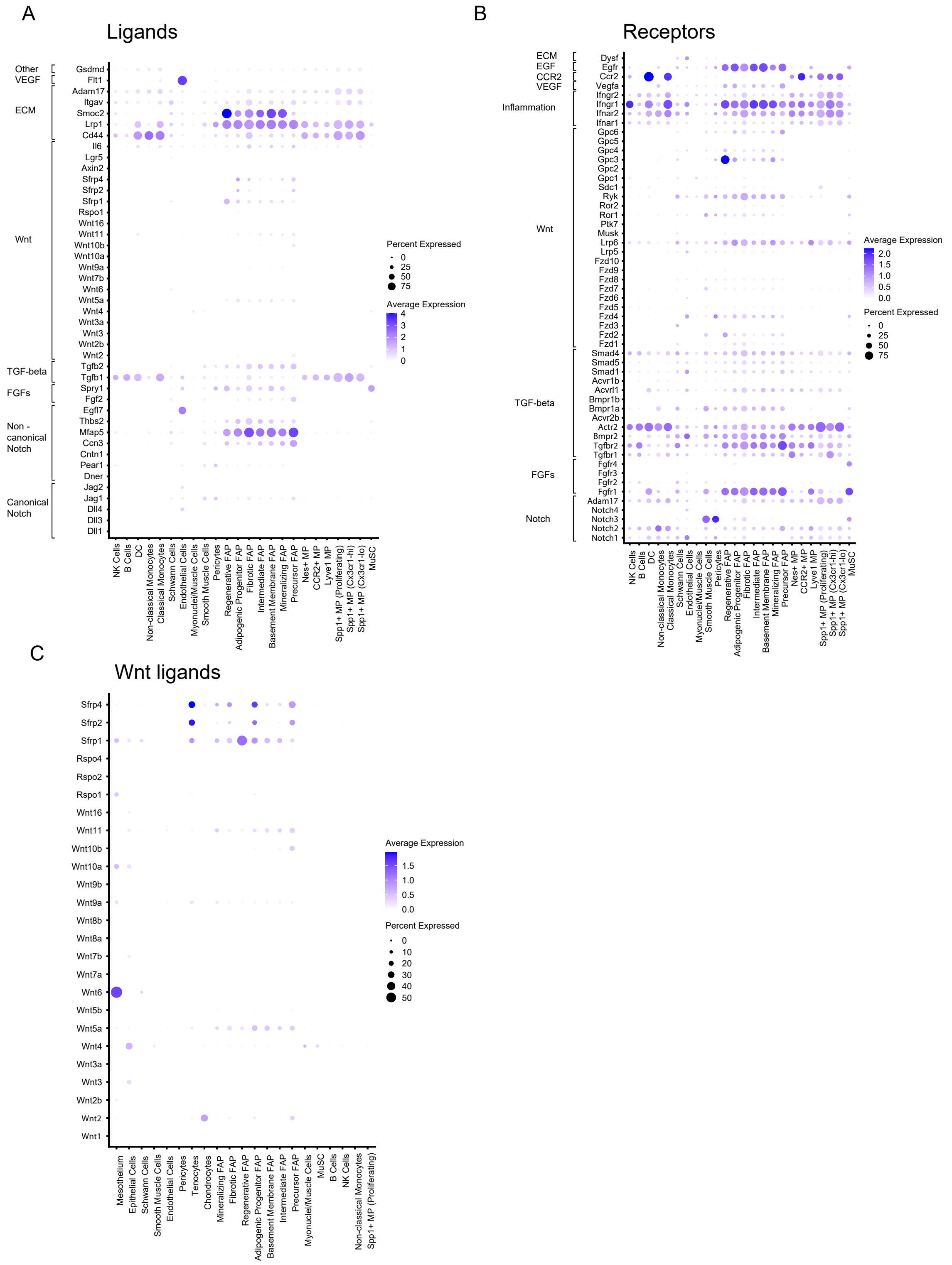


**Figure S8. Expression of ligands and receptors causally implicated in satellite cell proliferation after injury.** Refers to Figure 4.

(A) Dot plots indicating expression of ligands implicated in satellite cell proliferation and muscle regeneration after injury in cell populations resolved from single cell RNA-sequencing.
(B) Dot plots representing expression of receptors implicated in satellite cell proliferation and muscle regeneration after injury in cell populations resolved from single cell RNA-sequencing.
(C) Dot plots indicating expression of Wnt ligands in cell populations resolved from single cell RNA-sequencing

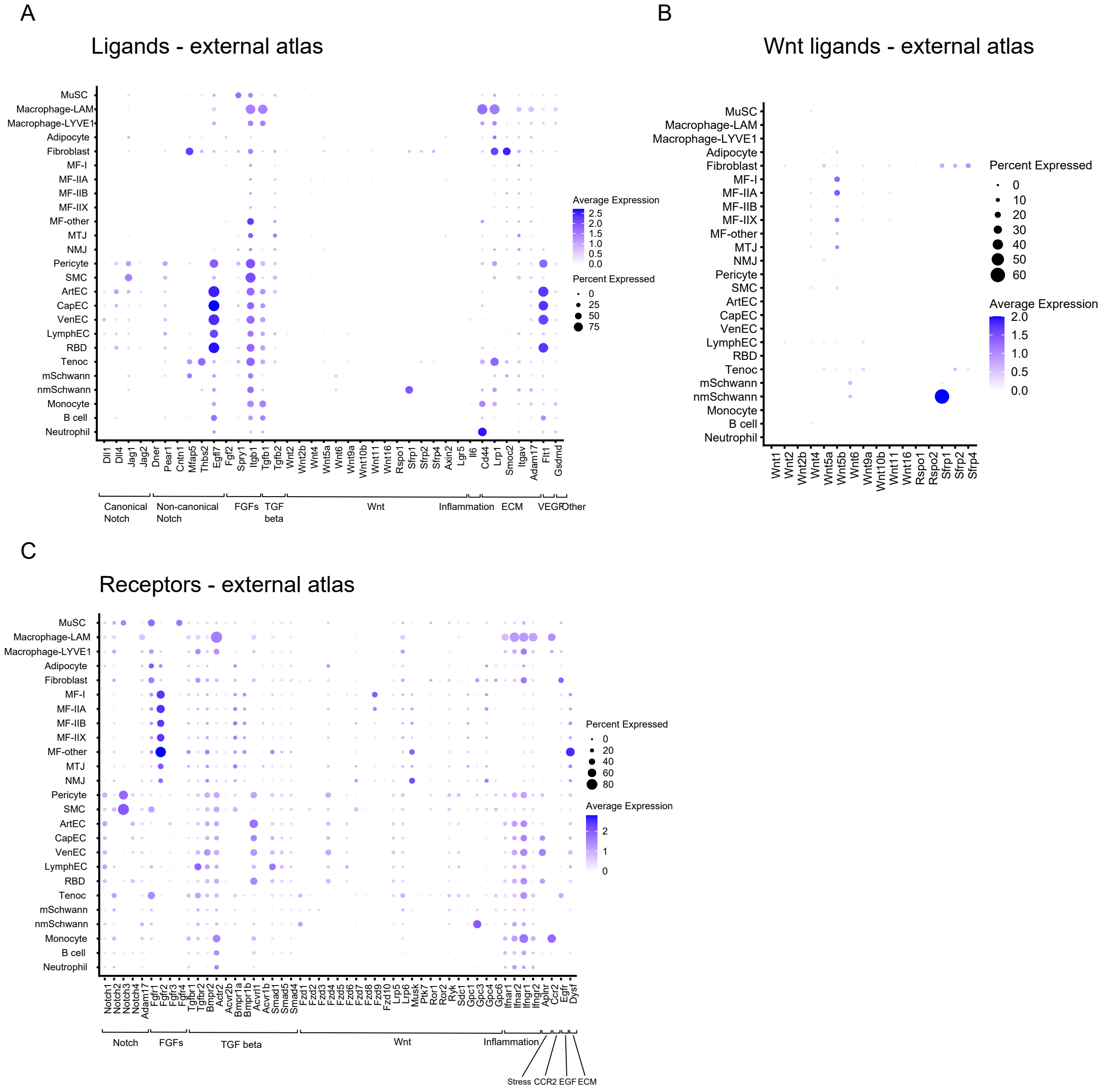


**Figure S9. Expression of ligands and receptors causally implicated in satellite cell proliferation after injury in a published single cell RNA-sequencing atlas of the skeletal muscle.**

(A) Dot plots indicating expression of ligands implicated in satellite cell proliferation and muscle regeneration after injury in cell populations resolved from single cell RNA-sequencing.
(B) Dot plots representing expression of receptors implicated in satellite cell proliferation and muscle regeneration after injury in cell populations resolved from single cell RNA-sequencing.
(C) Dot plots indicating expression of Wnt ligands in cell populations resolved from single cell RNA-sequencing.

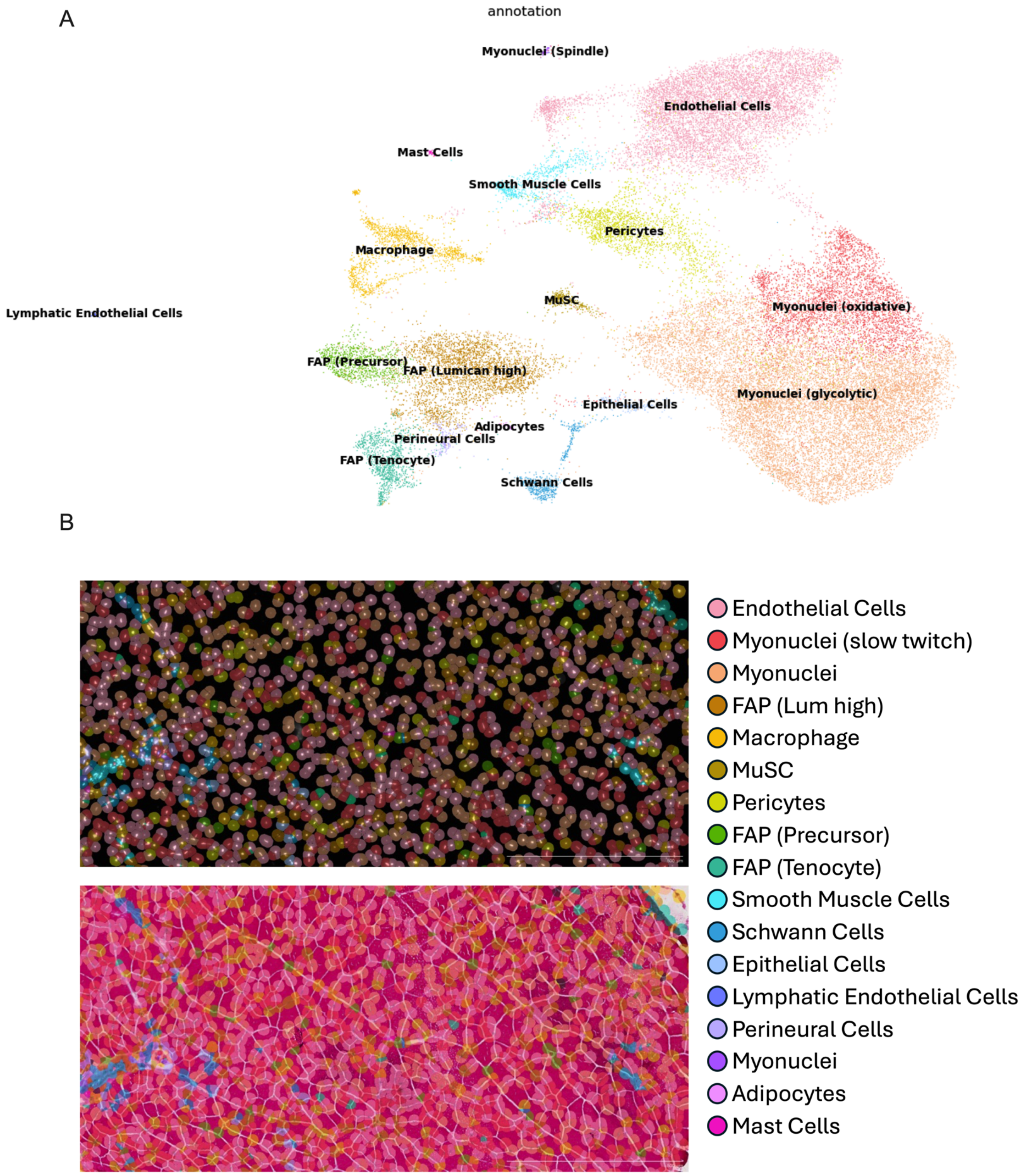

A
annotation
Myonuclei (Spindle)
Endothelial Cells
Mast Cells
Smooth Muscle Cells
Pericytes
Macrophage
MuSC
Myonuclei (oxidative)
Lymphatic Endothelial Cells
FAP (Precursor)
FAP (Lumican high)
Epithelial Cells
Myonuclei (glycolytic)
Adipocytes
Perineural Cells
FAP (Tenocyte)
Schwann Cells
B
Endothelial Cells
Myonuclei (slow twitch)
Myonuclei
FAP (Lum high)
Macrophage
MuSC
Pericytes
FAP (Precursor)
FAP (Tenocyte)
Smooth Muscle Cells
Schwann Cells
Epithelial Cells
Lymphatic Endothelial Cells
Perineural Cells
Myonuclei
Adipocytes
Mast Cells

**Figure S10. Spatial transcriptomic analysis of skeletal muscle in young adult and old mice 10 days after influenza A infection.**

(A) Cell clustering was performed on gene expression from Xenium in situ system images of mouse tibialis anterior muscle sections obtained with the Mouse Multi-Tissue standard panel using a nuclear segmentation algorithm. Cell clusters were annotated using marker gene expression. Cells were obtained from images of three sections on the same slide: young adult naïve mouse, a young adult mouse 6 days after influenza A infection and an old mouse 6 days after influenza A infection.

(B) Spatial representation of resolved cell populations from (A) alone (top) and projected onto hematoxylin and eosin staining of the same muscle section (bottom) from a young adult mouse 10 days after infection with influenza A. See also Figure 4E and Supplemental Table 1. Scale bars 500 μm. Object can be explored at https://www.nupulmonary.org/. Images are a single section from one animal each
